# Handling difficult cryo-ET samples: A study with primary neurons from *Drosophila melanogaster*

**DOI:** 10.1101/2023.07.10.548468

**Authors:** Joseph Y. Kim, Jie E. Yang, Josephine W. Mitchell, Lauren A. English, Sihui Z. Yang, Tanner Tenpas, Erik W. Dent, Jill Wildonger, Elizabeth R. Wright

## Abstract

Cellular neurobiology has benefited from recent advances in the field of cryo-electron tomography (cryo-ET). Numerous structural and ultrastructural insights have been obtained from plunge-frozen primary neurons cultured on electron microscopy grids. With most primary neurons been derived from rodent sources, we sought to expand the breadth of sample availability by using primary neurons derived from 3^rd^ instar *Drosophila melanogaster* larval brains. Ultrastructural abnormalities were encountered while establishing this model system for cryo-ET, which were exemplified by excessive membrane blebbing and cellular fragmentation. To optimize neuronal samples, we integrated substrate selection, micropatterning, montage data collection, and chemical fixation. Efforts to address difficulties in establishing *Drosophila* neurons for future cryo-ET studies in cellular neurobiology also provided insights that future practitioners can use when attempting to establish other cell-based model systems.

## INTRODUCTION

Cryo-electron tomography (cryo-ET) is a cryo-electron microscopy (cryo-EM) sub-discipline that has provided extraordinary ultrastructural insights of macromolecules, viruses, organelles, and cells in three-dimensions (3D) at high-resolution (Hylton & Swulius, 2021). With adherent cells, this can be achieved by culturing cells onto electron microscopy (EM) grids similarly to culturing cells on glass substrates (Hampton *et al*., 2017). The specimens can then be plunge-frozen in a cryogen close to liquid nitrogen temperatures to preserve them in a near-native state for cryo-imaging in a transmission electron microscope (TEM) without the need for chemical fixation, dehydration, plastic resin embedding, or heavy metal staining, which introduce artifacts and potentially alter cellular ultrastructure (Resch *et al*., 2011, Korogod *et al*., 2015). To collect data for cryo-ET, projection images are recorded on film or a camera while the specimen is tilted across a range of angles producing a ‘tilt series’. The images and the angular information of the tilt series are then computationally combined to produce a back-projection of the original 3D volume (Oikonomou & Jensen, 2017). Such volumes, or tomograms, provide immediate 3D information of the sample at nanometer-level resolution. Higher resolution structures of isolated complexes within a 3D tomogram are produced by averaging thousands of macromolecules of interest through a process described as sub-tomogram averaging (STA), thereby unlocking structural biology *in situ* (Wan & Briggs, 2016).

Cellular neurobiology has undergone a structural biology renaissance due to cryo-ET of cultured primary neurons, induced-pluripotent stem cells (iPSCs), and organoids (Bauerlein *et al*., 2017, Wu *et al*., 2023, Hoffmann *et al*., 2021). Novel insights include the detection and possible identification of luminal particles inside microtubules, the gradient of fascin-associated actin to cofilin-associated actin in neuronal growth cone filopodia, and mesophasic organization of putative GABA_A_ receptors in inhibitory synapses (Garvalov *et al*., 2006, Hylton *et al*., 2022, Liu *et al*., 2020). Using neurons for cryo-ET studies also comes with the additional advantage in that the 200-250 nm thick cellular extensions or neurites (e.g., axons or dendrites) that they produce to communicate intercellularly are amenable for direct cryo-TEM imaging without requiring thinning techniques, such as cryo-FIB-milling (Zuber & Lucic, 2022, Pesaresi *et al*., 2015, Chereau *et al*., 2017, Windebank *et al*., 1985). These thin neurites contain many common cytoskeletal proteins and organelles that are found in other cells, such as microtubules, actin, mitochondria, and endoplasmic reticulum and associated vesicles (Foster *et al*., 2022).

Many of the cryo-ET studies of primary neurons have been from mammalian sources, such as rodents or human iPSCs (Hylton *et al*., 2021, Wu et al., 2023). While such sources provide close relevancy for human health, assessments of neurons from multiple phyla can provide useful comparison points. A study that used cryo-ET and respiration assays to compare purified mitochondria ultrastructure between mouse liver, mouse heart, and entire fruit flies revealed potential correlation between inner mitochondrial organization and aging, and how such correlation was much more pronounced for less complex metazoans such as fruit flies (Brandt *et al*., 2017). While cells, including neurons, across different species may share common organelles, how those organelles are organized and operated is likely to be different when comparing these species, details of which may be revealed by cryo-ET investigations.

Challenges remain to fully realize the potential of cryo-ET for cell biology and, specifically, cellular neurobiology. Preparing primary neurons for cryo-ET is difficult when compared to simpler cells, such as bacteria. Even amongst adherent cell cultures, culturing primary neurons is challenging for a myriad of reasons. First, mature neurons do not undergo cell division, which automatically limits the supply of cells for starting material (Gordon *et al*., 2013). Genetic manipulation of primary neurons is also difficult because neurons are not as amenable to common viral transduction and calcium phosphate transfection methods (Sariyer, 2013). Neurons extend long neurites from the soma, which when cultured on glass confers a wide span of data acquisition by light microscopy imaging modalities. However, when neurons are cultured on EM grids, these extensions along with the cell body may physically limit themselves to the metal grid bars versus over the transparent, thin carbon film support. This effectively limits sample accessibility for modes of TEM data collection. In addition, the thinness and length-spans of neurites when compared with other cellular structures, may lend them to being more fragile during the processes of grid blotting and plunge-freezing. Axonal varicosities, defined as enlarged, oval-shaped swellings along narrow neurite shafts, have been noticeably present in most cryo-ET studies done on primary neurons (Foster et al., 2022, Schrod *et al*., 2018, Ma *et al*., 2022, Wu et al., 2023, Liu et al., 2020, Tao *et al*., 2018, Fischer *et al*., 2018). When such swellings appear *in vivo* or *in vitro*, they have been associated with neuronal damage, such as neurodegeneration or mild traumatic brain injury (Sun *et al*., 2022a). It was also demonstrated that puffs of buffer at low pressure were enough to induce varicosity formation in cultured neurons, suggesting that it may be possible that processes of blotting and plunge-freezing could induce varicosities due to contact pressure (Gu *et al*., 2017, Ma et al., 2022). Whether neurite varicosities in cryo-EM and cryo-ET studies are physiologically relevant or are artifacts of sample preparation is unknown.

Owing to the delicate nature of neurons and their sensitivity to being cultured on EM grids, additional forms of stabilization may be required prior to plunge-freezing. Conventional biological electron microscopy of cellular samples relies on chemical fixation with glutaraldehyde and paraformaldehyde first before osmication, dehydration, and resin embedding. In some cases, chemical fixation has also been combined with high-pressure freezing to preserve the ultrastructure of sensitive cell regions, such as nodes of Ranvier (Sosinsky *et al*., 2005). Single-particle cryo-EM can also use mild chemical fixation techniques to preserve structural integrity of individual macromolecules or to crosslink fragile multi-protein complexes (Kastner *et al*., 2008, Shukla *et al*., 2014). Chemical fixation prior to plunge-freezing has been used in cryo-ET for a variety of reasons, such as the inactivation of viruses, cell morphology time course analyses, or to stabilize fragile bacterial membrane protrusions (Klein *et al*., 2020, Mageswaran *et al*., 2021, Subramanian *et al*., 2018). However, when chemical fixation is applied, careful assessments must be made to determine whether artifacts have been introduced that could create observable differences between fixed and unfixed samples.

Here we present the use of primary neurons derived from 3^rd^ instar larvae of *Drosophila melanogaster* and adaptations to culture methods for cryo-ET imaging studies (Egger *et al*., 2013, Lu *et al*., 2015, Voelzmann & Sanchez-Soriano, 2022, Sibert *et al*., 2021). We highlight challenges associated with cryo-ET investigations of neurons, the steps taken to mitigate damage to cells and cellular ultrastructure, and to control varicosity frequency. We used grids with fewer holes in the thin film support, EM grid micropatterning, and chemical fixation of the cells before freezing. We were able to improve neuron integrity based on structural examination of neurites and the visible reduction in varicosities, yet these effects were not completely suppressed with the fixation protocols employed. This study highlights use of *Drosophila melanogaster* as a primary neuron culture platform for cryo-ET and discusses challenges and optimization of parameters used in preparing neurons for cryo-imaging.

## RESULTS

We first established a pipeline to prepare primary neurons from *Drosophila melanogaster* for cryo-ET studies (Fig 1). We used a *Drosophila* strain whose neurons were fluorescently labeled with a membrane-tethered green fluorescent protein (mCD8:GFP) that illuminated all neuronal membranes (Figure 2A and 2E) or a strain that expressed fluorescent markers to distinguish between dendrites and axons (Table 1). We used 3^rd^ instar larvae because they are at peak developmental size with many neurons (Tennessen & Thummel, 2011), and were easy to handle and dissect. Brains were extracted and underwent a series of digestions, spins, and washes to disrupt the tissue and homogenize it to a point where individual cells could be plated onto EM grids coated with concanavalin A. After 3-4 days of culturing with insect cell culture medium, neurites were seen extending along the grid squares. After culturing and light microscopy inspection of the grids, they were plunge-frozen for cryo-ET studies.

**Figure 1.**
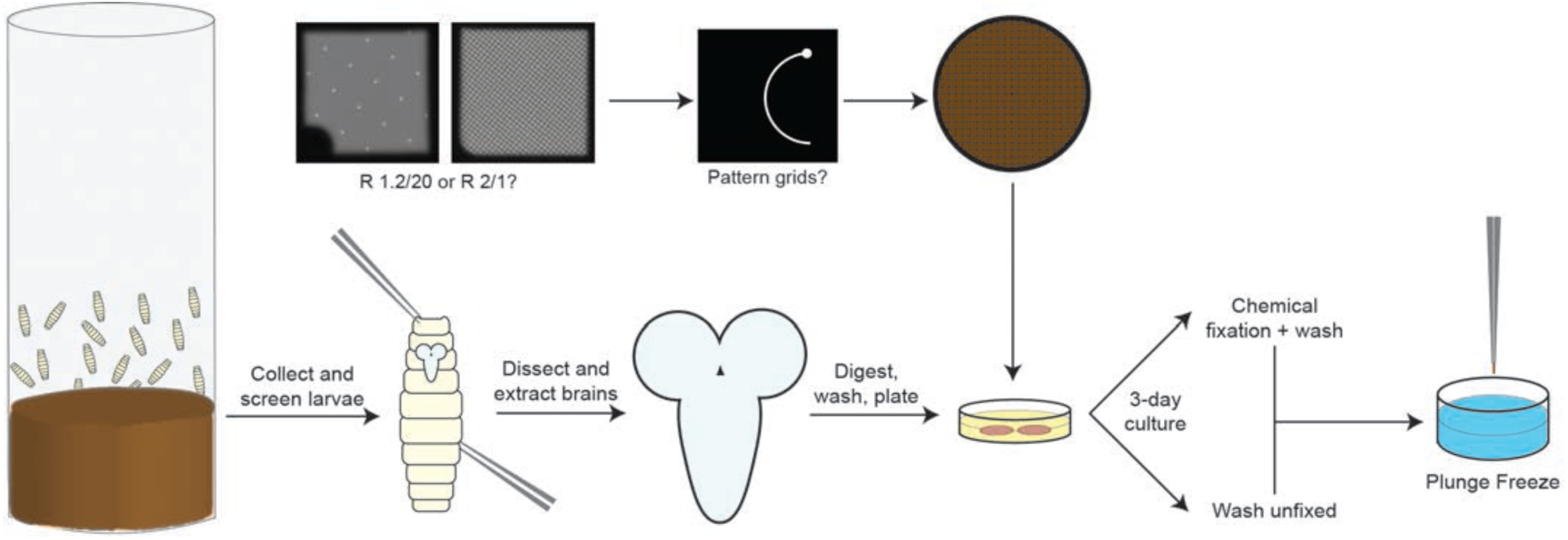
General workflow to culture primary neurons from *Drosophila melanogaster* for cryo-ET sample preparation. Approximately 30-40 larvae in the 3^rd^ instar developmental stage are picked from the food vial, easily identified by larvae crawling up the walls of the vial. The larvae are then screened, washed, sterilized, and then dissected under a dissecting microscope in room-temperature dissection saline to extract the brain with sharp tweezers. The brains are then digested to cellular suspensions, washed, and plated onto pre-coated EM grids incubating in media. The EM grids can have different hole sizes and spacing, or they can be micropatterned. The neurons are grown for at least 3 days, where neurites can be seen emerging from the cell body, and then plunge-frozen for cryo-ET. The neurons can either be washed without any alterations or treated with drugs and/or chemical fixatives before washing and freezing.

**Figure 2.**
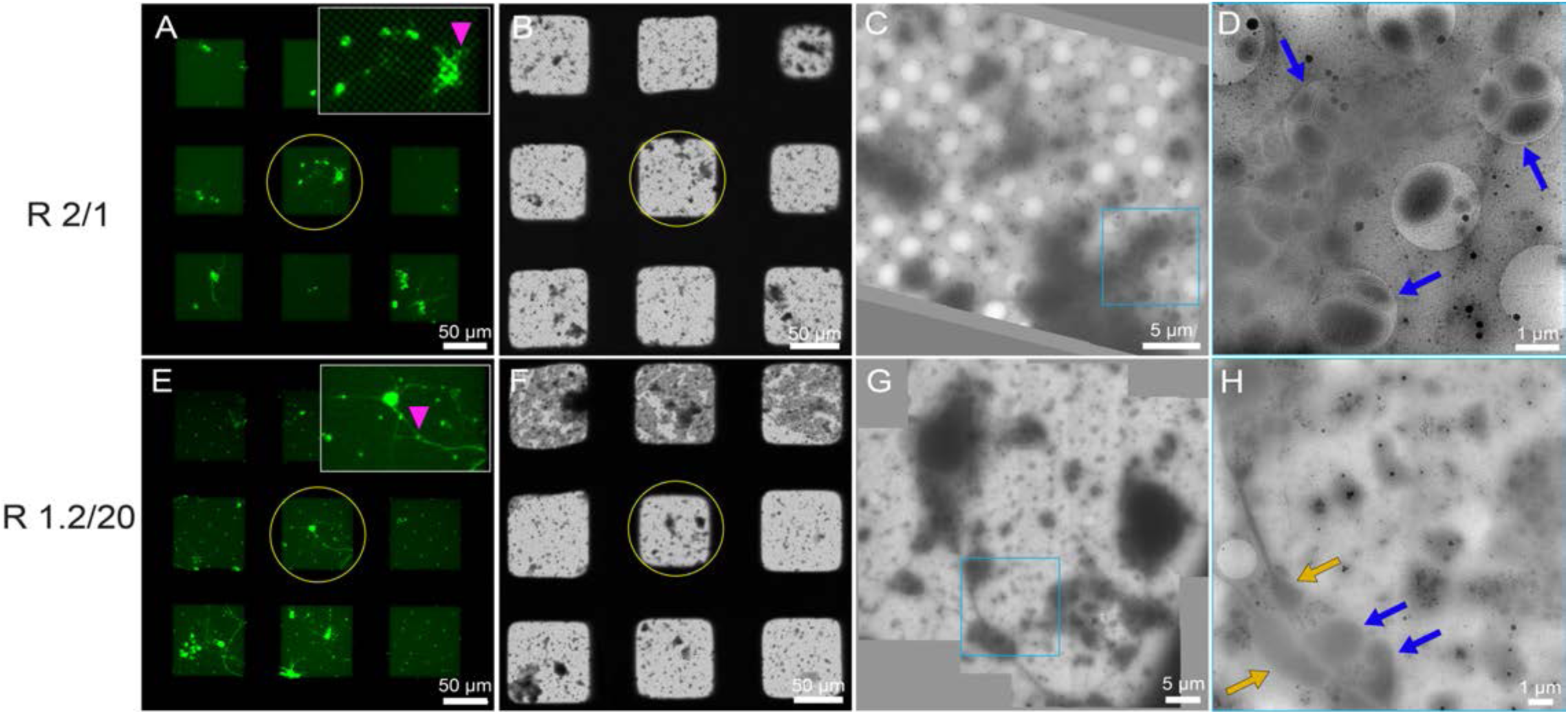
Initial attempts of cryo-EM/cryo-ET of primary neurons derived from *Drosophila melanogaster* larvae, grown for 4 DIV. **(A)** Live-cell fluorescent microscopy (FLM) image of *Drosophila* neurons expressing membrane-targeted GFP on unpatterned EM grid squares with fluorescent concanavalin A. The grid substrate has 2 µm holes spaced 1 µm apart (R 2/1). Inset is a zoomed-in image of the central neuron, with an intact neurite marked by the pink arrowhead. Green: *Drosophila* neurons. **(B)** Cryo-EM grid montage of the same grid in (A) after plunge-freezing on the Leica EM GP. Yellow circle show the same grid square. **(C)** Magnified cryo-EM square montage of the yellow circle in (A) and (B) maps **(D)** Magnified view of the blue square in (C), where the cellular region has no neurite linearity and is instead replaced by membrane blebs (blue arrow). **(E)** Live-cell fluorescent microscopy (FLM) image of *Drosophila* neurons expressing membrane-targeted GFP on unpatterned EM grid squares with fluorescent concanavalin A. The grid substrate has 1.2 µm holes spaced 20 µm apart (R 1.2/20). Inset is a zoomed-in image of the central neuron, with an intact neurite marked by the pink arrowhead. Green: *Drosophila* neurons. **(F)** Cryo-EM grid montage of the same grid in (E) after plunge-freezing on the Leica EM GP. Yellow circle show the same grid square. **(G)** Magnified cryo-EM square montage of the yellow circle in (E) and (F) maps. **(H)** Magnified view of the blue square in (G), where the cellular region has little neurite linearity and is instead surrounded by membrane blebs (blue arrow). The remaining intact neurites also contain ovoid swellings with heterogeneous diameters along its neurite path, known as varicosities (orange arrow). The scale bars in (A)-(H) are embedded in the image. Pixel sizes for (B) and (F) are 1,101 Å/pixel (82x magnification), pixel size for (C),(D),(G), and (H) are 17.12 Å/pixel (4,800x magnification).

**Table 1.** The *Drosophila* strains that were used in this study. While the mCD8::GFP strain was the primary strain used for the majority of this work, the Tau::GFP; DenMark strain was used as well, with no differences seen between the two at the cryo-imaging level.

| Strain | Fluorescent Marker 1 | Fluorescent Marker 2 | Screening | Purpose |
| --- | --- | --- | --- | --- |
| mCD8::GFP | Membrane-tethered GFP | N/A | Positive larvae have brains with GFP signal | Ubiquitous FLM confirmation of neuronal cells |
| Tau::GFP; DenMark | GFP-labeled microtubules (Axon-specific) | Membrane-tethered mCherry (Dendrite marker) | Negative larvae are short and squat ('Tubby') | Distinguish between axons and dendrites |

Upon cryo-EM imaging, we noticed alterations to the cells that were not identifiable at the light microscopy level. When neurons were cultured on grids with either 2 µm holes spaced 1 µm apart (R 2/1) or 1.2 µm holes spaced 20 µm (R 1.2/20) apart, swellings were spotted along the neurite (Figure 2). These swellings resembled the varicosities that have been reported in past studies (Ma et al., 2022, Fischer et al., 2018, Schrod et al., 2018). We noted that at both the neurite and cell body, membrane blebs, defined as spherical, blister-like membrane protrusions commonly seen in processes of apoptosis and cytotoxicity, were frequently seen surrounding the periphery, (Ponuwei, 2017, Lesort *et al*., 1997). These membrane blebs were often seen adjacent to or even replaced varicosity-enriched neurites. Comparisons with fluorescence light microscopy (FLM) images taken on cells before plunge-freezing and cryo-EM images of the same region confirmed that the neurites were often replaced with discrete membrane blebs (Figure 2). We hypothesized that the *Drosophila* neurons may become damaged between FLM imaging and cryo-EM imaging, including a progression from varicosities to a complete loss of neurite integrity.

The remnants of *Drosophila* neurons present on either R 2/1 or R 1.2/20 gold 200 mesh grids were not suited for cryo-ET data collection. However, the neurons had been healthy and had grown well on the grids when inspected at the live-cell FLM level (Figure 2A and E). For cryo-preservation, the grids were blotted from the backside using a Leica EM GP, similar to prior studies (Resch et al., 2011). The disruption of *Drosophila* neurons was very different when compared to mouse neurons, which grew easily on R 2/1 gold 200 mesh grids and remained intact after plunge-freezing, albeit with varicosities noted along neurites (Supplementary Figure 1). We screened *Drosophila* neurons to differentiate between intact neurites or damaged neurites that were engulfed with blebs. Cryo-ET data could not be acquired of intact *Drosophila* neurons until we micropatterned the R 2/1 grids prior to neuron culturing (Sibert et al., 2021). The condition of neurons and neurites further improved when *Drosophila* neurons were cultured on micropatterned R 1.2/20 grids. We collected multiple 3×3 and 3×4 montage tilt series using the recently developed Montage Parallel-Array Cryo-Tomography (MPACT) method (Yang *et al*., 2022) to capture intact neurites. Tilt series reconstruction, filtering, and segmentation of these tomograms revealed the presence of organelles, such as the endoplasmic reticulum (ER), mitochondria, microtubules, and vesicles (Figure 3).

**Figure 3.**
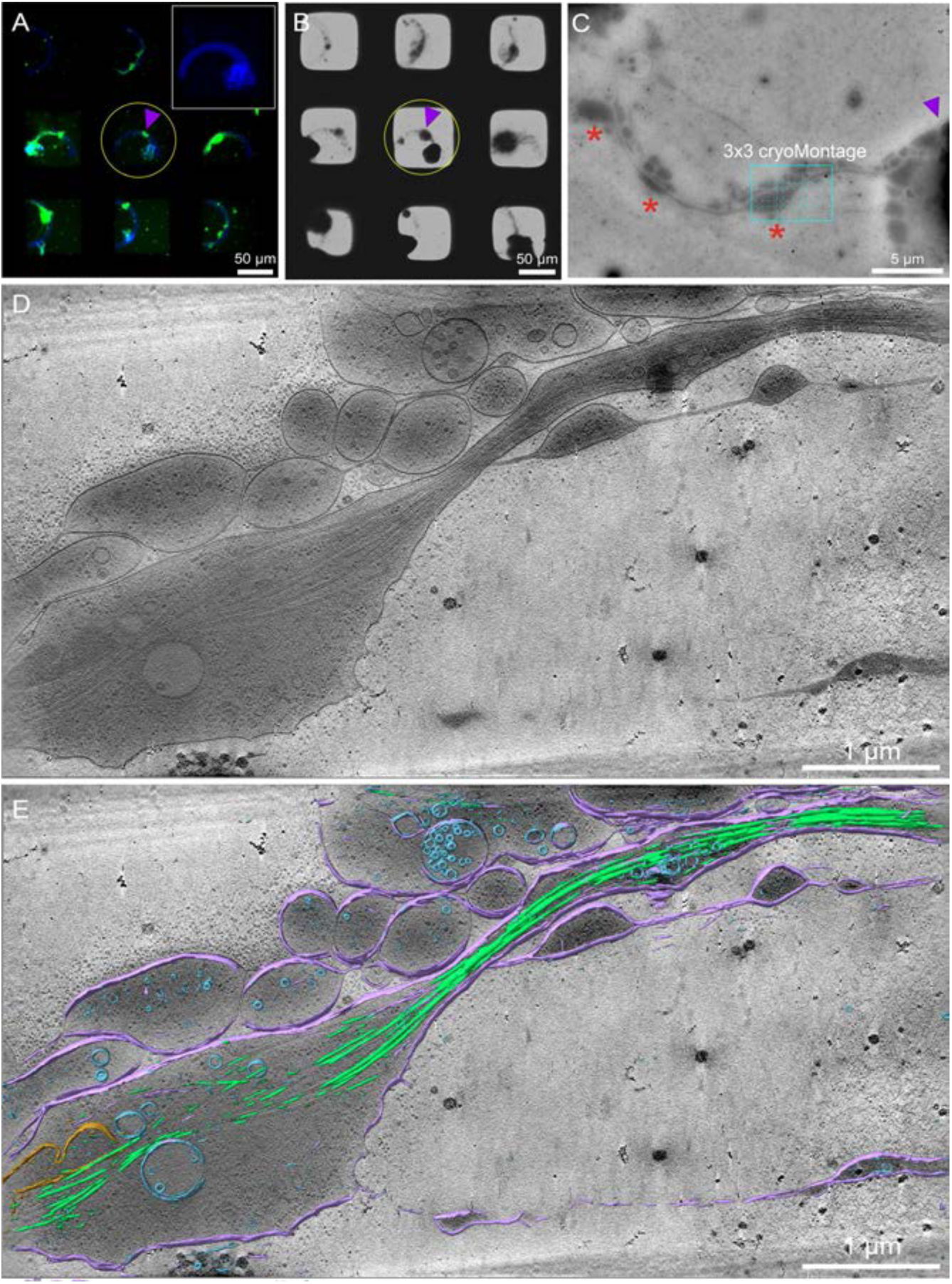
Maskless photopatterning and cryo-ET of primary neurons derived from *Drosophila melanogaster* larvae, grown for 3 DIV. **(A)** Live-cell fluorescent microscopy (FLM) grid montage of *Drosophila* neurons expressing membrane-targeted GFP on micropatterned R 1.2/20 grid squares with fluorescent concanavalin A. A curved pattern was used to increase available cellular space for data acquisition by cryo-ET on the thin neurites. The inset showcases the fluorescently-labeled micropattern used within the target grid square at higher contrast (yellow circle). The cell body is marked by the purple arrowhead. Green: *Drosophila* neurons. Blue: Photopattern **(B)** Cryo-EM grid montage of the same grid in (A) after plunge-freezing on the Leica EM GP. Yellow circle shows the same grid square from (A). **(C)** Magnified cryo-EM square montage of the yellow circle in (A) and (B). maps. Varicosities along the neurite path are represented by the red asterisks. **(D)** A 50 nm thick slice from a 3×3 montage tilt series that was taken at the cyan-marked box in (C) **(E)** Segmentation of the 3×3 montage tomogram by a convolutional neural network in EMAN2, overlaid on top of the 50 nm slice. Peripheral membranes and membrane blebs are colored in purple, microtubules are colored in green, vesicles are colored in blue, and mitochondria is colored in orange. The scale bars in (A-E) are embedded in the image. Pixel sizes for (B) is 1,101 Å/pixel (82x magnification), 24.1 Å/pixel (3,600x magnification) for (C), and 4.603 Å/pixel (19,500x magnification) for (D) and (E).

Based on the live-cell FLM images with good neurite integrity versus the cryo-EM images with damaged neurites (Figure 2), we sought to determine which sample handling and preservation manipulation may have impacted cellular morphology. Specifically, we wondered if the plunge-freezing process had the most impact because it involves direct physical contact and pressure on the substrate and cells during the filter paper-driven blotting process. Past reports have indicated that cellular damage from blotting could occur on fish keratinocytes, *Drosophila* S2 cells, *Plasmodium berghei* sporozoites, and Vero cells (Small *et al*., 2008, Lepper *et al*., 2010, Urban *et al*., 2010). To investigate this further, we took live-cell FLM images of the grid pre-blot and post-blot (Figure 4A and B). Post-blot images were obtained after blotting but without plunge-freezing the grid by using the ‘test-blot’ feature on the Leica EM GP. After blotting, the grid was returned to the culturing dish for re-imaging. Comparisons of pre-blot and post-blot FLM images of the same grid square show striking differences. Primarily, the membrane delineations on both the cell body and the neurites that the GFP signal highlights were reduced and lost after blotting (Fig 4A and B). This occurred regardless of whether the neurons were cultured on unpatterned or micropatterned grids (Supplementary Figures 2 and 3). Such diffuse signal could be compared to the varicosities and membrane blebs that are seen in cryo-EM images, along with the loss of neurite integrity.

**Figure 4.**
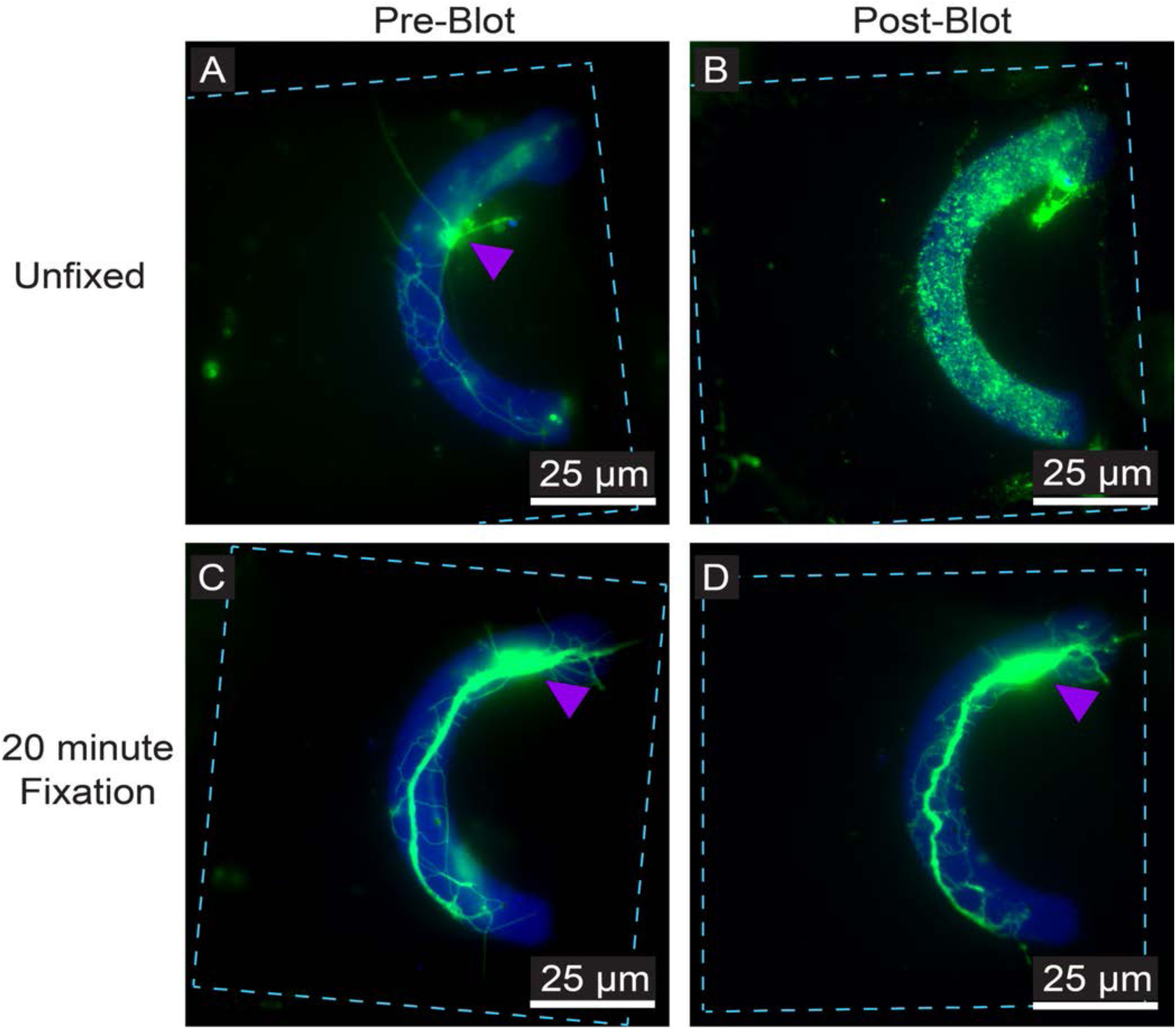
Effects of blotting on primary *Drosophila* neurons cultured on micropatterned EM grids under regular or chemically fixed conditions. **(A)** Live-cell fluorescent microscopy (FLM) of a *Drosophila* neuron expressing membrane-targeted GFP cultured on micropatterned R 1.2/20 grid squares with fluorescent concanavalin A. **(B)** The same square in (A), re-imaged after blotting the grid on the Leica EM GP. Note the complete loss of morphological integrity at both the cell body and the neurites, seen by diffuse green fluorescence scattered throughout the pattern. **(C)** FLM image of a *Drosophila* neuron expressing membrane-targeted GFP cultured on micropatterned R 1.2/20 grid squares with fluorescent concanavalin A. **(D)** The same square in (C), re-imaged after chemically fixing the grid in 4% paraformaldehyde in Kreb’s buffer and sucrose (PKS) for 20 minutes and then blotting the grid on the EM GP. Unlike (B), a vast majority of the cell remains intact after blotting. Grid bars are represented by the cyan dashed line. Cell bodies are marked by the purple arrowheads. The scale bars in (A-D) are embedded in the image. Green: *Drosophila* neurons. Blue: Micropattern.

To test whether the neuronal varicosities were induced by the blotting, we used chemical fixation combined with cryo-preservation. Chemical fixatives, such as paraformaldehyde or glutaraldehyde, crosslink covalent bonds across the biological sample thereby artificially preserving cell morphology (Eltoum *et al*., 2001). We questioned whether the crosslinking agents would provide added rigidity and greater mechanical strength to the *Drosophila* neurons, consequentially reducing varicosity formation and potential cellular damage. Live-cell FLM images were taken of grid squares with cells on them, and the grids were then fixed in a solution of 4% paraformaldehyde with Kreb’s buffer and sucrose (PKS) for up to 20 minutes (Dent & Meiri, 1992). The cells were then washed with 1X phosphate buffered saline (PBS) to remove all traces of the PKS, and the cells underwent the same blotting test on the EM GP as the unfixed cells did before imaging. Comparisons between post-blot unfixed and fixed neurons showed stark differences in cell morphology. The fixed neurons were well-preserved and retained intact neurites and cell bodies (Figure 4C and D). While cell preservation was not perfect, as seen by undulation and occasional loss of neurites between the pre-blot and the post-blot images, it was considerably better compared to unfixed neurons. Like the unfixed neurons, this effect occurred regardless of whether the neurons were cultured on unpatterned or micropatterned grids (Supplementary Figures 4-6).

Cryo-EM images of fixed *Drosophila* neurons confirmed that chemical fixation better preserved neuron integrity after blotting and plunge-freezing. We noticed that by fixing the neurons before plunge-freezing, the neuron integrity is maintained throughout the entire pipeline without needing to micropattern the grids, something that was previously a very rare occurrence. We observed reduced frequency of membrane blebs and varicosities even at a low magnification grid montage viewpoint, and fine neurites were easily observed that are amenable for high-throughput data collection (Figure 5A-C). Tomograms reconstructed from 3×3 tilt series from the MPACT scheme on these fixed neurites revealed that many macromolecules and organelles are still present within the cellular regions despite its chemical perturbation. Such features include mitochondria, vesicles, protein condensates, and actin filaments (Figure 5).

**Figure 5.**
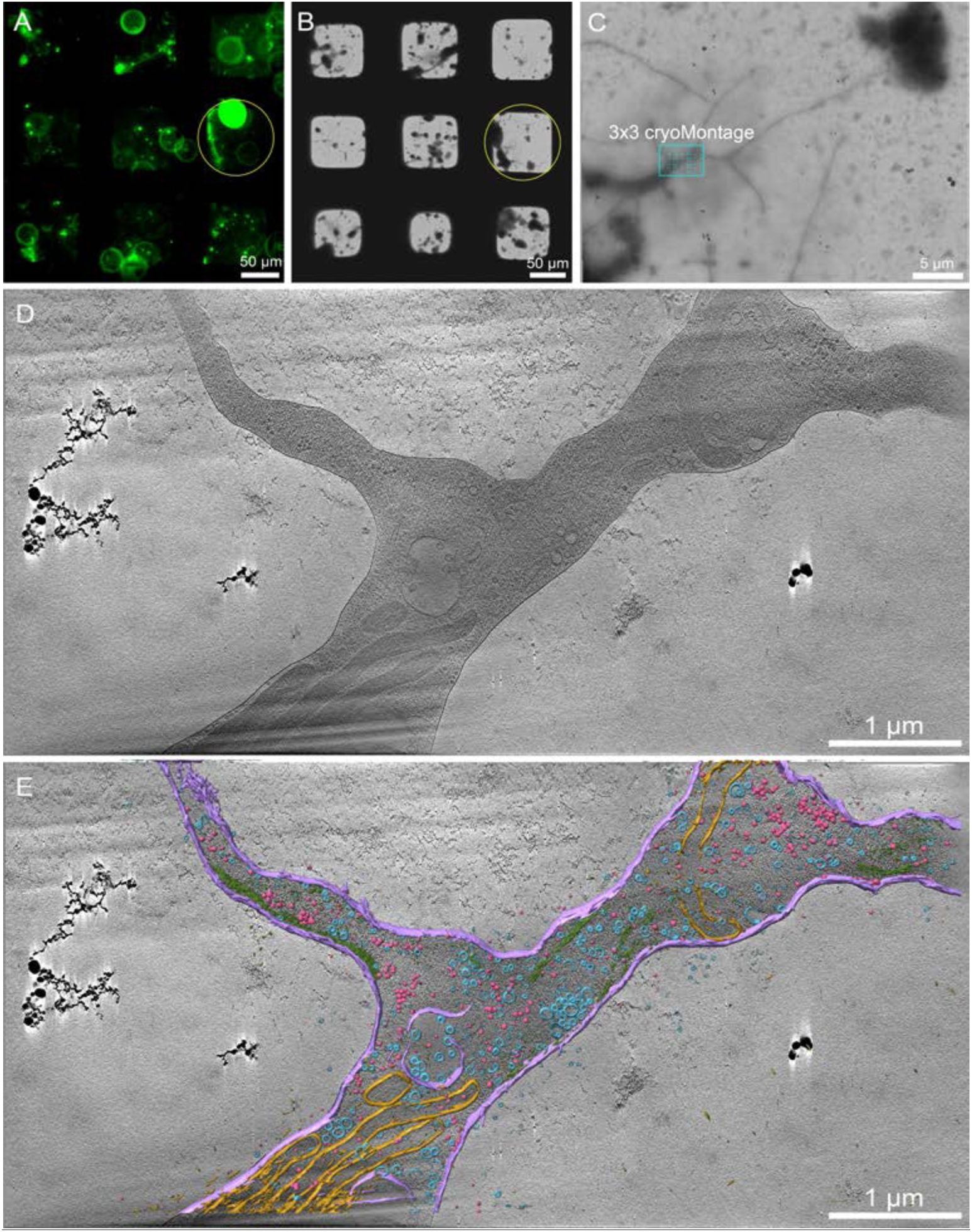
Cryo-ET of primary neurons derived from *Drosophila melanogaster* larvae, grown for 3 DIV and chemically fixed before freezing. **(A)** Live-cell fluorescence microscopy (FLM) grid montage of *Drosophila* neurons expressing membrane-targeted GFP on unpatterned R 1.2/20 grid squares coated with concanavalin A. Green: *Drosophila* neurons. **(B)** Cryo-EM grid montage of the same grid in (A) after chemical fixation in 4% PKS for 10 minutes and then plunge-freezing on the Leica EM GP. Yellow circle shows the same grid square. **(C)** Magnified cryo-EM square montage of the yellow circle in (A) and (B). **(D)** A 50 nm thick slice from a 3×3 montage tilt series that was taken at the cyan-marked box in (C) **(E)** Segmentation of the 3×3 montage tomogram by a convolutional neural network in EMAN2, overlaid on top of the 50 nm slice in (D). Peripheral membranes are colored in purple, macromolecular condensates are colored in pink, vesicles are colored in blue, mitochondria are colored in orange, and actin is colored in green. The scale bars in (A-E) are embedded in the image. Pixel sizes for (B) is 1,097 Å/pixel (82x magnification), 18.6 Å/pixel (4,800x magnification) for (C), and 4.518 Å/pixel (19,500x magnification) for (D) and (E).

With the observation that chemically fixed *Drosophila* neurons had considerably better cellular integrity and reduced frequency of varicosities, we decided to quantitatively examine the nature of these varicosities. We defined varicosities as oblong-shaped swellings that have diameters at least three times larger than the flanking neurites, based on past work that examined these objects present in tomograms of mouse neurons (Ma et al., 2022). Such swellings can become as large as 6 µm, which can be easily resolved by light microscopy yet were rarely seen by FLM before plunge-freezing. We measured the diameter of the varicosities (D_Varicosity_) at the median of the oblong-like swelling, and two sets of diameters at the ends of the varicosities where the membrane becomes parallel. The first set of diameters (D_Shaft1_, ‘Adjacent’) was measured and averaged at the ends of the varicosity, while the second set (D_Shaft2_, ‘Away’) was measured and averaged 250 nm away from where D_Shaft1_ was. D_Shaft2_ was measured to consider the possibility that the shaft diameters could expand or contract as it travels further from the varicosity. Measurements were taken across four different sample conditions (Table 2), using medium magnification cryo-EM maps of grid squares. Sample condition A consisted of *Drosophila* neurons cultured on curve-micropatterned R 1.2/20 grids that were frozen after chemical fixation with 4% PKS. Sample condition B consisted of *Drosophila* neurons cultured on curve-micropatterned R 1.2/20 grids that were frozen without chemical fixation. Sample condition C consisted of *Drosophila* neurons cultured on X-micropatterned R 2/1 grids that were frozen without chemical fixation. Sample condition D consisted of *Mus musculus* neurons cultured on unpatterned R 2/1 grids that were frozen without chemical fixation. We defined D_Varicosity_:D_Shaft_ ratio as a method to assess the varicosity while considering the thickness of the neurite. We utilized both D_Varicosity_ and D_Varicosity_:D_Shaft_ ratio to categorize the varicosities into four classes across all sample conditions: 1) high D_Varicosity_ and high D_Varicosity_:D_Shaft_, 2) low D_Varicosity_ and high D_Varicosity_:D_Shaft_, 3) high D_Varicosity_ and low D_Varicosity_:D_Shaft_, and 4) low D_Varicosity_ and low D_Varicosity_:D_Shaft_ (Figure 6). Despite the variety of sample conditions with respect to grid hole spacing, micropatterning, chemical treatment, or species, varicosities were observed in all four samples (Figure 6, Supplementary Figures 7-8).

**Figure 6.**
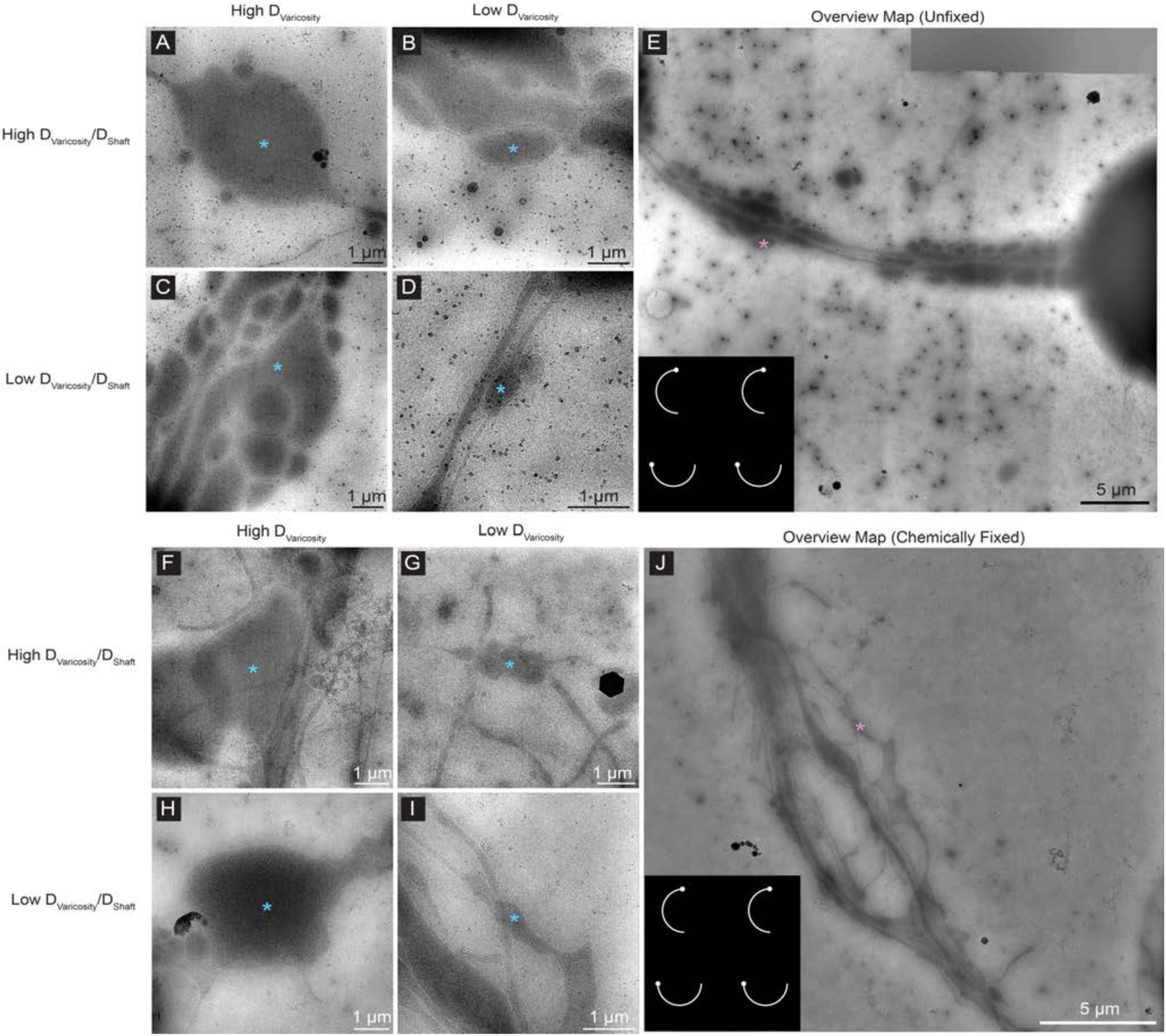
Categorization of varicosities based on the diameters of the varicosities (D_Varicosity_) and the ratio between D_Varicosity_ and D_Shaft_, (the diameter of the neurite shafts adjacent to the varicosity) in both unfixed and chemically fixed neurons. All varicosity classes (blue asterisk) and the representative overview map (A-E) were from unfixed *Drosophila* neurons on curve-patterned R 1.2/20 grids, while all varicosity classes (blue asterisk) and the representative overview map (F-J) were from chemically fixed *Drosophila* neurons on curve-patterned R 1.2/20 grids. **(A)** An unfixed varicosity class defined by having a high D_Varicosity_ and a high D_Varicosity_:D_Shaft_ ratio. **(B)** A class defined by having a low D_Varicosity_ and a high D_Varicosity_:D_Shaft_ ratio. **(C)** A class defined by having a high D_Varicosity_ and a low D_Varicosity_:D_Shaft_ ratio. **(D)** A class defined by having a low D_Varicosity_ and a low D_Varicosity_:D_Shaft_ ratio. **(E)** An overview cryo-EM map of a *Drosophila* neuron cultured on curve-patterned R 1.2/20 grids, plunge-frozen on the Leica EM GP without chemical fixation. The pattern used is shown in the inset, and the pink asterisk is the varicosity class in (B). **(F)** A chemically fixed varicosity class defined by having a high D_Varicosity_ and a high D_Varicosity_:D_Shaft_ ratio. **(G)** A class defined by having a low D_Varicosity_ and a high D_Varicosity_:D_Shaft_ ratio. **(H)** A class defined by having a high D_Varicosity_ and a low D_Varicosity_:D_Shaft_ ratio. **(I)** A class defined by having a low D_Varicosity_ and a low D_Varicosity_:D_Shaft_ ratio. **(J)** An overview cryo-EM map of a *Drosophila* neuron cultured on curve-patterned R 1.2/20 grids, plunge-frozen on the Leica EM GP after chemical fixation. The pattern used is shown in the inset, and the pink asterisk is the varicosity class in (I). The scale bars in (A-J) are embedded in the image. Pixel sizes are 15.3 Å/pixel (5,600x magnification) for (A-E), 13.7 Å/pixel (6,500x magnification) for (F) and (H-J), and 18.8 Å/pixel (4,800x magnification) for (G).

**Table 2.** The four different sample conditions that were tested for neurite varicosity measurements studies by cryo-EM.

| Sample Condition | Species | Hole spacing | Micropattern | Chemical fixation before freezing? |
| --- | --- | --- | --- | --- |
| A | *Drosophila melanogaster* | R 1.2/20 (1.2 µm holes spaced 20 µm apart) | Curved pattern | Yes, 4% PKS |
| B | *Drosophila melanogaster* | R 1.2/20 (1.2 µm holes spaced 20 µm apart) | Curved pattern | No |
| C | *Drosophila melanogaster* | R 2/1 (2 µm holes spaced 1 µm apart) | X-Pattern | No |
| D | *Mus musculus* | R 2/1 (2 µm holes spaced 1 µm apart) | Unpatterned | No |

We measured these varicosities and plotted them for all four sample conditions (Figure 7, Supplementary Figure 9). Of the four conditions, we note there was a significant reduction in the average D_Varicosity_:D_Shaft_ ratio, which ranged from a 31% reduction in the ‘Away’ ratio from condition B to A to a 49% reduction in the ‘Adjacent’ ratio from condition C to A) for the chemically fixed *Drosophila* neurons when compared to conditions without chemical fixation (Figure 7E-F, Table 2). Excluding the fixed sample condition A, it was observed that the average D_Varicosity_:D_Shaft_ ratios amongst the three unfixed samples were not heavily altered, though there was a slight variation in the ‘Adjacent’ D_Varicosity_:D_Shaft_ set of sample conditions (Figure 7E-F). This was in spite of the fact that D_Varicosity_, ‘Adjacent’ D_Shaft1_, and ‘Away’ D_Shaft2_ were quite varied across all four sample conditions (Supplementary Figure 9E-G). Scrutinizing the measurements more closely, we noted that the unfixed *Drosophila* neurons on curve-micropatterned R 1.2/20 grids (Sample condition B) had a larger average D_Varicosity_ while the D_Varicosity_ for the other three sample conditions were quite similar (Supplementary Figure 9E, Table 2). Upon examining the D_Shaft_ set of measurements for both ‘Adjacent’ and ‘Away’ measurements, we discovered that the D_Shaft_ of neurons under sample conditions A and B, which were cultured on R 1.2/20 grids, were significantly larger than the D_Shaft_ of neurons under sample conditions C and D, which were cultured on R 2/1 grids (Supplementary Figure 9F-G). Differences in species from which the neurons were derived did not make a difference in these measurements, as sample condition C was derived from *Drosophila* and D was derived from *Mus musculus*. Despite the variation in sample conditions and their corresponding measurements for D_Varicosity_, ‘Adjacent’ D_Shaft1_, and ‘Away’ D_Shaft2_, we noted the reduction in varicosities for chemically fixed samples to be the most significant. This indicated that there may be a correlation between the extent of varicosity size and neuronal damage during cryo-sample preparation.

**Figure 7.**
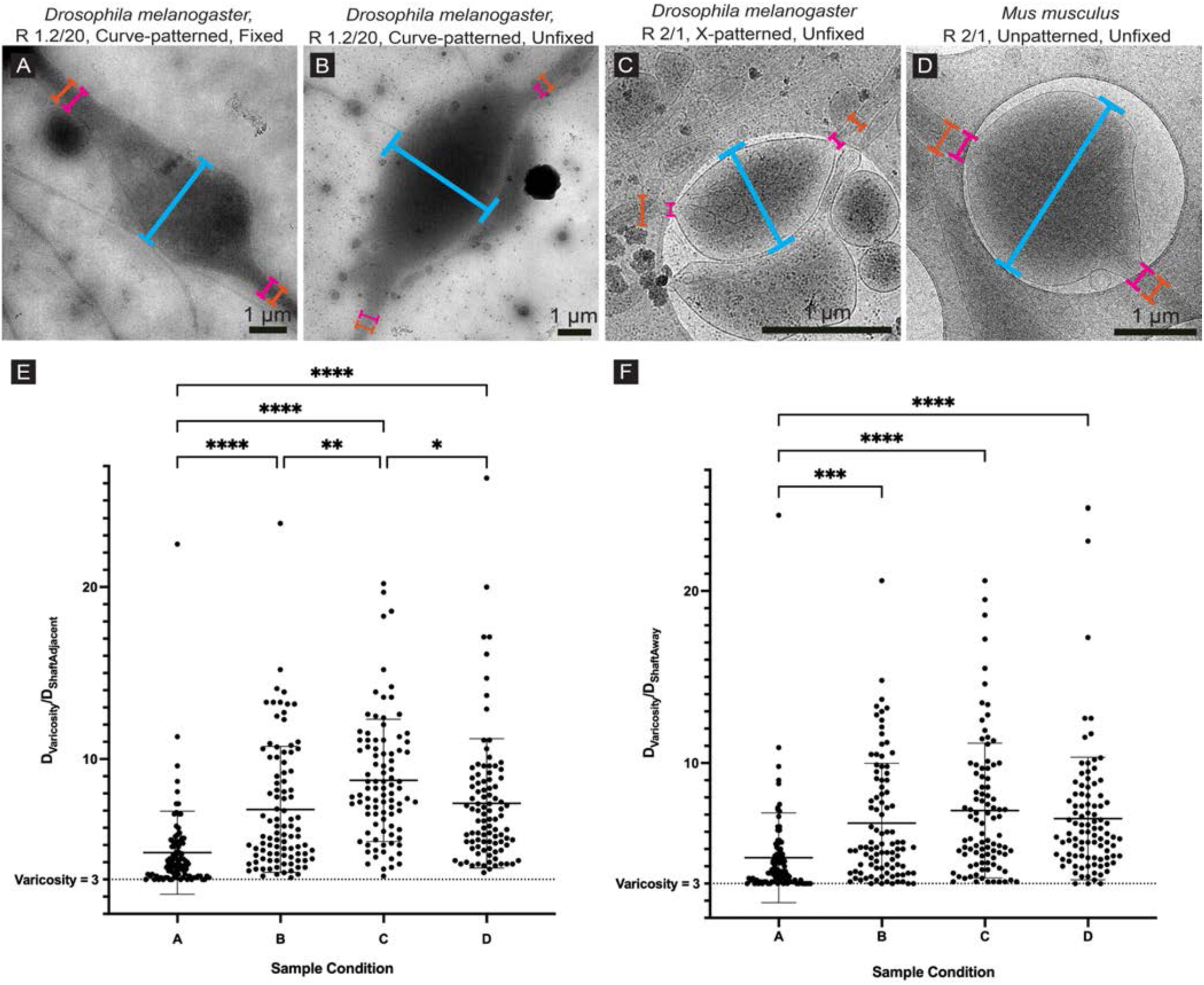
Comparisons of D_Varicosity_:D_Shaft_ ratios across different neuronal sample conditions, measured within overview cryo-EM maps. Two D_Varicosity_:D_Shaft_ ratio measurements are done for each sample condition due to varying D_Shaft_ with respect to its distance from the varicosity. D_Varicosity_ (Cyan in A-D) is measured at the median between the two ends, while D_Shaft1_ (Pink in A-D, or ‘Adjacent’ shaft) is measured at the two ends of the varicosity where the membranes are parallel and D_Shaft2_ (Orange in A-D, or ‘Away’ shaft) is measured 250 nm away from D_Shaft1_. **(A)** A representative varicosity from sample condition A, where Drosophila neurons were cultured on curve-patterned R 1.2/20 grids and plunge-frozen after chemical fixation. **(B)** A representative varicosity from sample condition B, where Drosophila neurons were cultured on curve-patterned R 1.2/20 grids and plunge-frozen without chemical fixation. **(C)** A representative varicosity from sample condition C, where *Drosophila* neurons were cultured on X-patterned R 2/1 grids and plunge-frozen without chemical fixation. **(D)** A representative varicosity from sample condition D, where *Mus musculus* neurons were cultured on unpatterned R 2/1 grids and plunge-frozen without chemical fixation. **(E)** Column scatter plot of D_Varicosity_:D_Shaft1_ ratios across the four sample conditions. Varicosities are defined by D_Varicosity_ being 3x the D_Shaft_, represented by the dashed line as a minimum threshold. **(F)** Column scatter plot of D_Varicosity_:D_Shaft2_ ratios across the four sample conditions. N = 96 varicosities for sample condition A, N = 95 for B, N = 93 for C, N = 95 for D. All data are mean ± s.d. * is P < 0.05, ** is P < 0.01, *** is P < 0.001, **** is P < 0.0001 calculated using a one-way ANOVA with post-hoc Tukey’s multiple comparison test. The sample conditions on the X-axis in (E-F) can be found in Table 1 and are also the same conditions as the respective panels (A-D) (i.e. Sample condition A in (E) is the same condition that is labeled in (A)).

Despite the reduction of varicosities, we noted that chemical fixation produced its own cellular artifacts. Most notably, microtubule integrity was compromised in samples that were chemically fixed compared to unfixed, varicosity-heavy neurons (Figure 8). In the majority of tomograms of chemically fixed *Drosophila* neurons, we failed to find microtubules, normally abundant and easy to identify by their characteristic thickness of approximately 25 nm and tubular shape (Foster et al., 2022, Atherton *et al*., 2018, Grange *et al*., 2017). Occasionally, microtubules were seen in fixed neurites; however, they were shrunken, frayed, and crooked (Figure 8C-D). Such observations were noted for all PKS-fixed neurites, and treatment with 5 or 20 nM of Taxol for 24 hours prior to PKS-fixation did not rescue this effect (Supplementary Figure 10A-D). Fixing the neurites with paraformaldehyde in cytoskeleton buffer (4% PCB) that is more amenable for tubulin stabilization also did not rescue the disappearance of microtubules from the neurites (Supplementary Figure 10E-F). Adding 0.25% glutaraldehyde (GA), a common EM fixative used to preserve cell ultrastructure and microtubules (Graham & Orenstein, 2007, Bartlett & Banker, 1984), to the 4% PKS partially rescued microtubule structure. However, many of the microtubules remained frayed and disrupted in appearance when compared to those present in unfixed neurites (Supplementary Figure 10G-H, Figure 8). An additional artifact that may have been caused by the fixative was a ‘clearing’ effect where patches within neurites appeared empty and contained no subcellular features, with sharp delineations between such patches and adjacent cellular regions (Figure 8C, Supplementary Figure 10C). This effect was more commonly observed in neurites fixed with 4% PKS + 0.25 GA (Supplementary Figure 10G). It is currently unknown if these artifacts are applicable to other cell cultures from different species, since subcellular and macromolecular composition is likely to vary across each sample type. To address the ‘clearing’ effect, while also rescuing microtubule structure, we increased the glutaraldehyde concentration and used other buffer systems, such as sodium cacodylate and sodium phosphate, commonly used for conventional EM studies (Ivanchenko *et al*., 2021, Grabrucker *et al*., 2009). A higher concentration of 2.5% glutaraldehyde mixed in 0.1 M sodium cacodylate buffer not only fully restored microtubule frequency, observed by the integrity of the tubular ultrastructure, protofilament striations, and visible tubulin dimers, but also greatly reduced the clearing artifacts seen in cells fixed with 4% PKS or 4% PKS + 0.25% GA (Figure 8E-F). Interestingly, fixing the cells with the same fixative, but with an addition of 2% paraformaldehyde seemed to partially disrupt the microtubule network; while most of the tubular morphology of the microtubules remained intact, slight fraying and breaks in the lattice were observed, though not to the same extent as cells fixed in 4% PKS (Supplementary Figure 10I-J).

**Figure 8.**
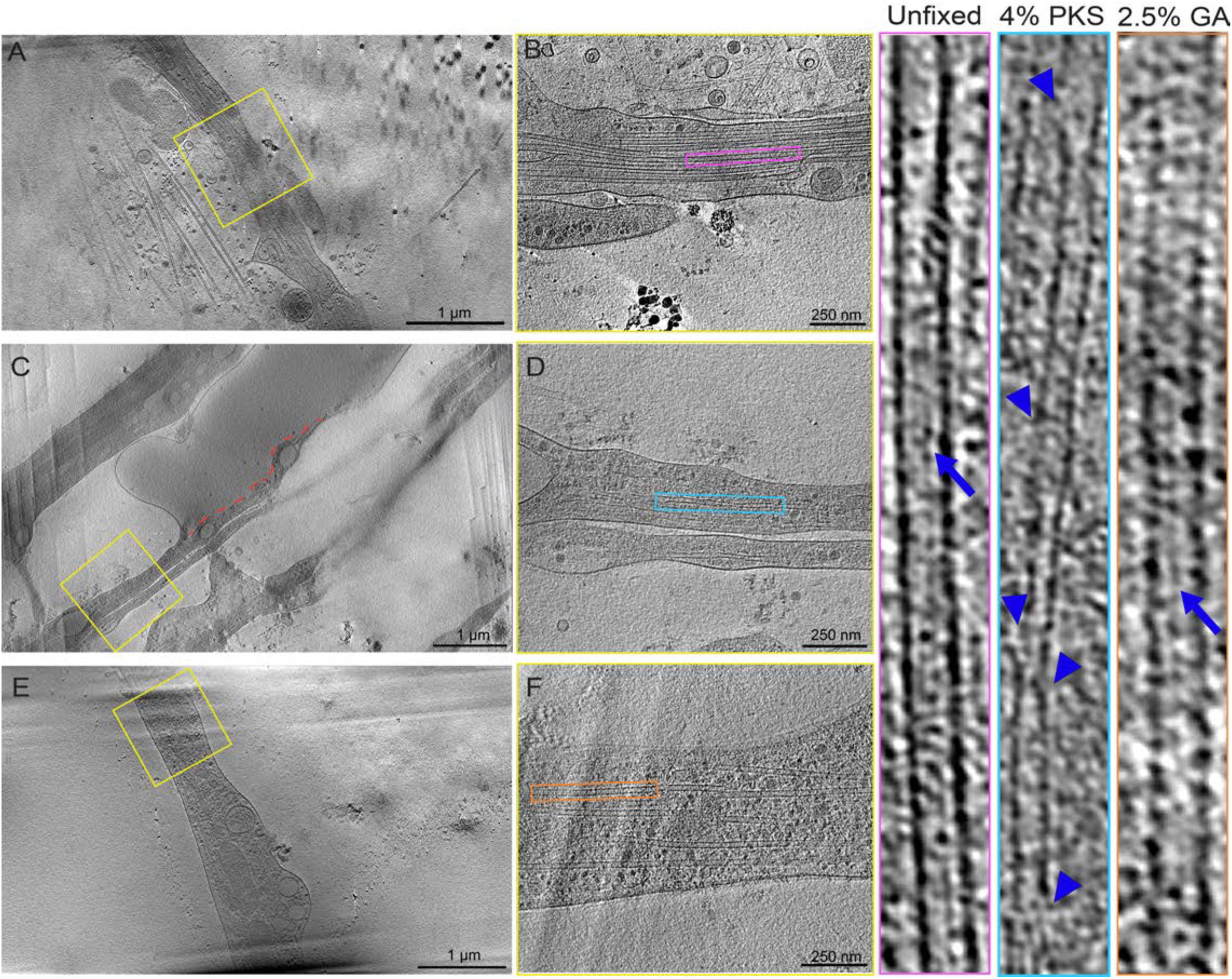
Comparison between tomograms of unfixed *Drosophila* neurons and fixed neurons shows that microtubules could be perturbed by chemical fixation, depending on the fixative used. **(A)** A 50 nm thick 3×3 montage tomogram slice on a *Drosophila* neuron cultured on curve-patterned R 1.2/20 grids, frozen without chemical fixation. **(B)** A zoomed up view of a 40 nm thick tomogram slice at the yellow box highlighted in (A). Bundles of intact microtubules can be seen. A higher magnification view of a single microtubule is shown on the right, marked by the pink box in B. Striations denoting individual microtubule protofilaments are marked by the blue arrow. **(C)** A 50 nm thick 3×4 montage tomogram slice on a *Drosophila* neuron cultured on curve-patterned R 1.2/20 grids, frozen after fixation in 4% paraformaldehyde in Krebs buffer supplemented with sucrose (PKS). The red dotted line depicts a possible artifact where there is a sharp boundary between regions containing subcellular density and empty, or ‘cleared’ areas **(D)** A zoomed up view of a 40 nm thick tomogram slice at the yellow box highlighted in (C). A higher magnification view of a single microtubule is shown on the right, marked by the cyan box in D. Compared to (B), the microtubule looks jagged and disrupted, with breaks along the protofilament marked by the blue arrowheads. **(E)** A 50 nm thick 3×3 montage tomogram slice on a *Drosophila* neuron cultured on curve-patterned R 1.2/20 grids, frozen after fixation in 2.5% glutaraldehyde (GA) in cacodylate buffer. **(F)** A zoomed up view of a 40 nm thick tomogram slice at the yellow box highlighted in (E). Unlike (C-D), bundles of intact microtubules can be seen in this fixed neurite sample. A higher magnification view of a single microtubule is shown on the right, marked by the orange box in F, where similarities can be made with the unfixed microtubule (pink), including striations denoting individual microtubule protofilaments marked by the blue arrow. Scale bars are embedded in the image. Pixel sizes for (A-D) are 4.603 Å/pixel (19,500x magnification) and 4.517 Å/pixel for (E-F).

We examined the ‘clearing’ artifacts more closely and wondered if this may be caused by imbalances in osmotic pressure while the neurons were fixed. Classical electron microscopy studies have shown that ultrastructural changes can occur in a variety of cells and tissues if the fixative has osmolarities/osmolalities that are non-isotonic (Hegele-Hartung *et al*., 1989, Bozzola & Russell, 1999). Therefore, we measured the osmolalities of all the solutions that the *Drosophila* neurons were cultured or incubated in using freezing point depression (Table 4). Through this, we discovered that both the 4% PKS fixative and the 4% PKS + 0.25% GA fixative was estimated to be substantially hypertonic by one order of magnitude when compared to 1X PBS and the media used to culture the cells. Strikingly, 2.5% GA in 0.1 M cacodylate buffer, the fixative that preserved microtubules the best while also reducing the clearing artifacts, was only slightly hypertonic compared to 1X PBS and the neuronal media. This suggested that the clearing artifacts that were frequently seen in neurons fixed in 4% PKS with or without 0.25% GA are from plasmolysis, effects that were ameliorated when the neurons were fixed in a solution that was closer to isotonic levels.

**Table 3.** Average varicosity diameters, average diameters of ‘Adjacent’ shafts and ‘Away’ shafts, and average ratio of varicosity to shaft diameter measurements for both ‘Adjacent’ shafts and ‘Away’ shafts with standard deviation, categorized by the four different sample conditions that were tested. N = 96 varicosities for condition A, 95 for condition B, 93 varicosities for condition C, and 95 varicosities for condition D.

| Sample Condition | Average $D_{\text{Varicosity}}$ (nm) | Average 'Adjacent' $D_{\text{Shaft1}}$ (nm) | Average 'Away' $D_{\text{Shaft2}}$ (nm) | Average of 'Adjacent' $D_{\text{Varicosity}}:D_{\text{Shaft1}}$ ratios | Average of 'Away' $D_{\text{Varicosity}}:D_{\text{Shaft2}}$ ratios |
| --- | --- | --- | --- | --- | --- |
| A | 1011 ± 672 | 233 ± 143 | 238 ± 144 | 4.6 ± 2.4 | 4.5 ± 2.6 |
| B | 1608 ± 1060 | 249 ± 153 | 270 ± 159 | 7.1 ± 3.6 | 6.5 ± 3.5 |
| C | 832 ± 349 | 108 ± 61 | 142 ± 82 | 8.8 ± 3.5 | 7.2 ± 3.9 |
| D | 960 ± 417 | 142 ± 59 | 156 ± 64 | 7.4 ± 3.7 | 6.8 ± 3.5 |

**Table 4.** Osmolality for all the solutions that were used during neuronal culturing, fixation, and wash steps before plunge-freezing. Raw osmolality values were measured by freezing point depression, then corrected with a calibration curve based on potassium chloride standards with known osmolalities for the final values. Osmolality values marked with ‘*’ are rough estimates, as the measurements for those samples was too high for the calibration curve to accurately correct.

| Sample | Osmolality<br>milliosmoles/kg<br>H<sub>2</sub>O (mOsm) |
| --- | --- |
| 1X Phosphate-Buffered Saline | 314 |
| 0.1 M Cacodylate Buffer | 222 |
| 0.1 M Phosphate Buffer | 222 |
| Supplemented Schneider's Media | 314 |
| 2.5% Glutaraldehyde in 0.1 M Cacodylate Buffer | 418 |
| 2.5% Glutaraldehyde in 0.1 M Phosphate Buffer | 423 |
| 2% Paraformaldehyde + 2.5% Glutaraldehyde in 0.1 M Cacodylate Buffer | 997 |
| 2% Paraformaldehyde + 2.5% Glutaraldehyde in 0.1 M Phosphate Buffer | 1009 |
| 4% Paraformaldehyde in Kreb's Buffer | 1447 |
| 4% Paraformaldehyde in Kreb's Buffer + Sucrose | **2037\*** |
| 4% Paraformaldehyde + 0.25% Glutaraldehyde in Kreb's Buffer + Sucrose | **1973\*** |

Despite the artifacts induced by chemical fixation, we observed organelles and macromolecules in fixed neurons that were comparable to unfixed neurons (Figure 9). We were able to find 40-60 nm diameter synaptic vesicles (Figure 9A) and larger, electron-dense 70-120 nm dense core vesicles (Figure 9H) in both samples (Tao et al., 2018, Foster et al., 2022). Longitudinal actin was identified (Figure 9B), as individual filaments or in packed bundles. Organelles and macromolecules for cellular biogenesis were also detected for both sets of samples and included ribosomes and the endoplasmic reticulum along neurite shafts (Figure 9C and E) (Nedozralova *et al*., 2022). The large presence of ribosomes within the neurites suggested that these cultured neurons were immature because ribosome numbers reduce dramatically when the axons mature and form synapses (Costa *et al*., 2019). Seen amongst both unfixed and fixed neurons were clusters and dispersions of biomolecular condensates (Figure 9G and 5D and E), membrane-less organelles involved in a wide variety of functions (Banani *et al*., 2017). A variety of mitochondrial morphologies were seen in both unfixed and fixed neurons, with variations in length, cristae organization, and fusion/fission dynamics (Figure 9D, Figure 3D and E, Figure 5D and E). We also observed cases of mitochondrial clustering in single neurite segments (Figure 5D and E), something that would not be resolvable by conventional light microscopy. Lastly, multivesicular bodies were seen (Figure 9F, Figure 5D and E), which demonstrated that chemical fixation could be used to arrest cells at certain time points, which would be useful for neurodegenerative studies where the autophagy-lysosomal pathway is disrupted (Riera-Tur *et al*., 2022).

**Figure 9.**
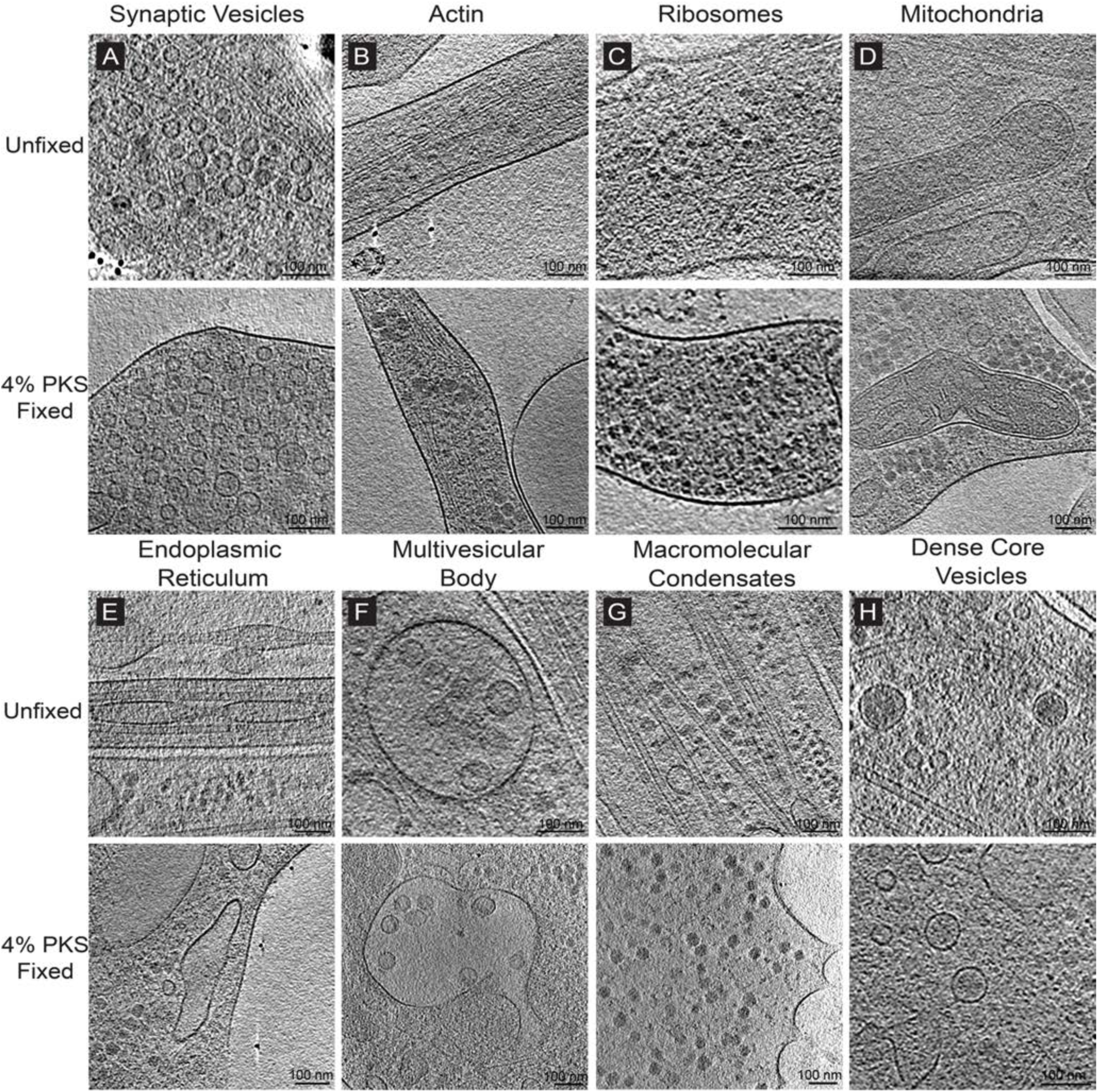
Comparison between tomograms of unfixed *Drosophila* neurons and fixed neurons shows that many well-known macromolecules and organelles remain intact after fixation with 4% PKS, despite perturbation of the microtubule network. **(A)** Comparison of synaptic vesicles in 50 nm tomogram slices of unfixed neurons (Top) and fixed neurons (Bottom) **(B)** Comparison of actin filaments in 25 nm tomogram slices of unfixed neurons (Top) and fixed neurons (Bottom) **(C)** Comparison of 25 nm ribosomes in tomogram slices of unfixed neurons (Top) and fixed neurons (Bottom) **(D)** Comparison of a mitochondria in 50 nm tomogram slices of unfixed neurons (Top) and fixed neurons (Bottom). **(E)** Comparison of the endoplasmic reticulum in 50 nm tomogram slices of unfixed neurons (Top) and fixed neurons (Bottom). **(F)** Comparison of a multivesicular body in 50 nm tomogram slices of unfixed neurons (Top) and fixed neurons (Bottom, also seen in Figure 5D-E). **(G)** Comparison of macromolecular condensates in 50 nm tomogram slices of unfixed neurons (Top) and fixed neurons (Bottom). **(H)** Comparison of dense core vesicles in 25 nm tomogram slices of unfixed neurons (Top) and fixed neurons (Bottom). Scale bars are embedded in the image. All tomogram slices in this figure are 25 or 50 nm thick and are derived from 3×3 or 3×4 montage tomograms. Pixel sizes for (A-D) are 4.603 Å/pixel (19,500x magnification) for the unfixed tomograms in (A-H) and 4.518 Å/pixel (19,500x magnification) for the fixed tomograms in (A-H).

## DISCUSSION

We began this study to establish *Drosophila melanogaster* as a model system for cryo-ET studies of primary neurons (Figure 1) (Kim *et al*., 2020). Substantial discoveries in neurobiology can be attributed to the *Drosophila* model system, including the discovery of ion channels involved with repolarization, chemosensation and somatosensation, genes and proteins involved with the circadian rhythm, and components of the cell signaling Notch pathway (Bellen *et al*., 2010). *Drosophila* neurons could be a complementary model to study cellular neurobiology by cryo-ET as well as an evolutionary comparison versus other primary neuronal cell models, such as embryonic neurons from rodent brains or from human iPSCs (Wu et al., 2023). An added advantage of utilizing *Drosophila* neurons was the ease and variety of genetic manipulations that were available, something we took advantage of by using a transgenic fly strain that expresses a pan-neuronal membrane-tethered GFP reporter for our primary analysis (Duffy, 2002, Lee & Luo, 2001, Koushika *et al*., 1996). A recent study attempted to analyze the intracellular compartments of neurites propagating from *Drosophila* neurons grown on EM grids (Foster et al., 2022). However, analysis was limited due to difficulties in sample handling for cryo-ET, difficulties which we also encountered and highlight here. We believe such insights will not only inform practitioners of cryo-ET on primary neurons, but also describe difficulties and potential pitfalls that could help a user who is trying to do any variant of cellular cryo-ET.

While initial culturing attempts were successful on EM grids at the live-cell FLM level with optimized handling and techniques, obtaining tilt series on neurites of good quality after plunge-freezing was not feasible. Throughout most of the square maps that we screened; we consistently saw broken neurites with swollen varicosities interspersed throughout the intact regions (Figure 2). Membrane blebs often accompanied these varicosities, and sometimes even replaced the neurites themselves. While varicosities are a feature commonly observed throughout the majority of past cryo-ET neuronal studies (Foster et al., 2022, Schrod et al., 2018, Ma et al., 2022, Wu et al., 2023, Liu et al., 2020, Tao et al., 2018, Fischer et al., 2018), the high levels of cell death and fragmentation that were seen prevented us from moving forward with data collection. This occurred regardless of hole spacing, as we did not see a substantial improvement in sample quality when the neurons were cultured on R 1.2/20 grids. Similar to a previous publication (Foster et al., 2022), we believe that *Drosophila* neurons are a particularly fragile sample for cryo-ET, undergoing stress and cell death from the grid manipulation during sample preparation. This was supported by the little difficulty we encountered culturing and collecting data on *Mus musculus* cortical neurons on R 2/1 grids (Supplementary Figure 1).

It was not until we micropatterned our grids that we started seeing more success in obtaining tilt series, as we were able to acquire data at intact cellular regions (Figure 3). Such regions can be identified as neurites emanating from the cell body in a continuous manner without loss of integrity and fragmentation into membrane blebs. Although we continued seeing varicosities and surrounding membrane blebs sporadically, sample quality improved considerably compared to *Drosophila* neurons cultured on unpatterned grids. MPACT tilt series data collection validated that within these varicosities, intracellular features were seen at high resolution that suggest healthy neurite growth (Figure 3D and E). It is not completely known why micropatterning would help *Drosophila* neurons survive plunge-freezing, but one hypothesis could be that the restricted microenvironment could fasciculate neurites together as they grow across the grid squares. A previous cryo-ET study of axonal varicosities from mouse cortical and hippocampal neurons found that bundles of axons grown on lacey carbon grids had significantly reduced varicosity frequencies compared to singular axons (Ma et al., 2022). Bundled axons make up the nerves that make up the central and peripheral nervous system, which showcases its physiological relevance and necessity (Sun *et al*., 2022b). If the varicosities here represent an intermediate morphology between healthy neurites and fragmented neurites, this could mean that forcing the neurites to bundle together by micropatterning could improve sample preparation for cryo-ET, though more studies will be needed to verify this.

By analyzing cell morphology at the light and cryo-EM/ET level and controlling the sample preparation process by substrate selection, micropatterning, and chemical fixation, we theorize that the pressure induced by blotting before plunge-freezing is a particularly sensitive step for cells cultured on grids for cryo-ET (Figures 3-5, Supplementary Figures 2-6). The physics and impact of sample blotting in cryo-EM has been studied and discussed for single-particle cryo-EM, and is believed to contribute to preferential orientation and denaturation at the air-water interface (Glaeser & Han, 2017, Glaeser, 2018, Armstrong *et al*., 2020). It is not too improbable to consider that blotting could decrease sample quality at the cellular level as well and has in fact been reported on in the past (Urban et al., 2010, Small et al., 2008, Lepper et al., 2010). For neurons, blotting appears to induce or accelerate the formation of varicosities that may lead to cell death, likely due to the pressure from the blotting process. The blotting pressure that is imposed by the Vitrobot was measured to be 20 kPa, (or 20 nN/µm^2^), or 150 mmHg, the equivalent of the systolic blood pressure of a patient with hypertension (Armstrong et al., 2020, Basile, 2002). A similar range of pressures is likely used for other instruments due to the overarching goal of producing a thin uniform layer of liquid in a few seconds. Such pressures could have an immediate impact on neurons; puffs of Hank’s buffer onto mouse hippocampal neurons at a pressure of ∼0.25 ± 0.06 nN/µm^2^ (kPa) were enough to induce varicosity formation (Gu et al., 2017). The type and force of pressure applied to a grid while blotted is not completely known, though shear stress has been proposed as a possible mechanism from capillary pressures (Armstrong et al., 2020). In addition, whether neurons on the grid experience the full pressure of the filter paper blotting or some downstream effect is also not well-understood. However, it is known that high-magnitude pressures can produce deleterious effects in neurons both *in vitro* and *in vivo*, such as neurite retraction in PC12 cells and tissue deformation in embryonic mesenchymal cells at pressures of 0.27 ± 0.4 nN/μm^2^ and 1.6 ± 0.8 nN/μm^2^ respectively (Voortman *et al*., 2014, Franze *et al*., 2009). When pressure is applied for a longer duration at greater magnitudes, this mechanical force can induce necrosis and apoptosis (Quan *et al*., 2014), which could explain the membrane blebs and loss of neurite integrity that were seen on initial cryo-imaging attempts (Figure 2).

Previous studies have shown that chemical fixation increases the stiffness of cell membranes. Cells fixed in 4% PFA had a Young’s Modulus measured by atomic force microscopy to be approximately five-fold higher than unfixed cells, while cells fixed in 0.5% GA had a Young’s Modulus measured to be approximately three-fold higher (Kim *et al*., 2017, Hutter *et al*., 2005). This higher stiffness could explain why chemically fixed neurons remained intact after blot testing or plunge-freezing (Figure 4-5, Supplementary Figures 4 and 6). Such stiffness is not only increased in fixed cells, but is also uniform across the cell surface, unlike unfixed cells where the membrane above the nucleus is a magnitude softer at 2-3 kPa than surfaces above regions where actin is prominent at 10-20 kPa (Yamane *et al*., 2000). Another possible reason fixed cells resisted blotting-induced damage could be due to the cross-linking provided by the chemical fixatives halting the activities of mechanosensitive (MS) channels on the neuronal cell surface. Touch-sensitive transient receptor potential (TRP) channels and the mechanosensitive ion channel Piezo1 are endogenously expressed on the surface of mouse hippocampal neurons, and blocking such receptors reduced puffing-induced varicosities (Shibasaki *et al*., 2007, Gu et al., 2017, Falleroni *et al*., 2022). *Drosophila melanogaster* also expresses mechanosensitive channels on larval sensory neurons, including pickpocket-1 and pickpocket-26, DmPiezo, and NOMPC, (Yan *et al*., 2013, Song *et al*., 2019, Adams *et al*., 1998, Mauthner *et al*., 2014, Gorczyca *et al*., 2014, Guo *et al*., 2014). Further studies will be needed to determine whether such channels are expressed on brain-derived larval neuronal cultures, and if selectively blocking them could reduce varicosity formation and cell death. It should also be noted that chemical fixation does not occur ubiquitously at equivalent rates across all cellular structures, as the presence of free amine groups is required for crosslinking. This is exemplified in how membrane blebbing was still observed in chemically fixed cells, as lipids tend to be unreactive to glutaraldehyde or paraformaldehyde due to their lack of free amino groups (Huebinger *et al*., 2018).

Despite testing a multitude of sample conditions including substrate selection, patterning, fixation, and even species, our analysis revealed that varicosities were seen in all conditions tested (Figure 6 and 7, Supplementary Figures 7-9). Early electron microscopy studies of fixed, resin-embedded sections from rat hippocampal slices identified varicosities across the CA1 to CA3 regions and defined them as axonal swellings whose diameter exceeded the diameter the adjacent axonal shafts by at least 50%. A large majority contained synaptic vesicles, indicating that varicosities can be functionally and physiologically relevant as synaptic components (Shepherd & Harris, 1998). However, it was also reported that such varicosities can be pathological from traumatic brain injury and neurodegeneration (Sun et al., 2022a, Gu et al., 2017, Nikic *et al*., 2011, Gao *et al*., 2011). Distinguishing between these two different forms of varicosities is important for defining the functional landscape of neurons for *in vitro* to *in vivo* experiments. With our results, we were able to disentangle the two different types of varicosities, indicating that the majority that remained are likely to be physiologically relevant and not trauma induced. This could be highlighted by the significant decrease in the extent of the swelling by our D_Varicosity_:D_Shaft_ ratio (Figure 7D-E), although we caution that this is not an exact correlation and that more studies are needed to explore this relationship further.

Despite the difficulty in sample preparation, tomograms from *Drosophila* neurons demonstrate that they could serve as a good model for *in situ* neuronal cryo-ET as a complement to primary neurons derived from rodents and hiPSCs (Figures 3, 5, and 9). Most organelles and macromolecules that are necessary for neuronal function were faithfully preserved between unfixed or fixed neurons. As neurites typically contain archetypal subcellular organelles and macromolecules but are thin enough to be imaged without FIB-milling, neurons are a unique model for cryo-ET studies where high-throughput data collection must be combined with cellular insights. Additional advantages include the plethora of transgenic *Drosophila* models that are available and the speed, ease, and low-cost of husbandry. Examples of technological innovation and new biological questions from *Drosophila* neurons include the development of the MPACT tilt series scheme (Yang et al., 2022), and raising a fly strain that can fluorescently distinguish between axons and dendrites for correlative light and electron microscopy (CLEM), the former which is now widely available for the cryo-ET community and the latter which is the subject of future studies.

Microtubule depolymerization proved to be an unexpected roadblock to implement chemical fixation into our cultured neuronal samples. The PKS fixation-induced depolymerization was persistent, resisting Taxol treatment and the use of a tubulin-friendly buffer (Figure 8, Supplementary Figure 10). Artifacts induced by chemical fixation have always been prevalent in electron microscopy and were one of the reasons as to why cryo-EM was developed. Studies comparing different imaging modalities have shown that chemical fixation by PFA can alter organelles and macromolecules from their native state, such as the ER and biomolecular condensates (Hoffman *et al*., 2020, Irgen-Gioro *et al*., 2022). While past studies have indicated that PFA fixation can disrupt dynamic microtubule networks in other cells and *in vitro* (Laporte *et al*., 2022, Gambarotto *et al*., 2021, Szasz *et al*., 1982), it was surprising that this occurred in neuronal microtubules, as they are believed to be stable compared to other microtubule networks due to their crucial roles in cellular architecture and axonal transport (Rolls *et al*., 2021, Baas *et al*., 2016). Neurons that were fixed by a mixture of PFA and GA, at proportions believed to preserve microtubules, still had artifacts intermediate or comparable to the rare microtubules that were seen in PKS-fixed cells (Supplementary Figure 10G-J). It was only when the cells were fixed with a higher concentration of GA without any PFA that microtubule integrity was restored (Figure 8E-F). While PFA-fixation is known to poorly preserve microtubules, little is known about how PFA disrupts microtubules at the ultrastructural level. GA seems to better preserve microtubule structure, but it is not known whether this is due to the chemical activity of GA itself, or differences in tonicity vs. PFA-solutions. It is also possible that buffers composition may also have an impact on microtubules, or a differences in the reaction rates of chemical cross-linking. Such questions may be addressed in future studies.

Patches where cellular content was missing after fixation (Figure 8C, Supplementary Figure 10C and E), or a disappearance of subcellular components (Wisse *et al*., 2010), represented another possible limitation of using chemical fixation before plunge-freezing. Based on osmolality measurements (Table 4), it seems that such patches could be due to excessive osmotic pressure from highly tonic solutions inducing plasmolysis. Chemical fixatives, especially paraformaldehyde, have a much higher osmotic pressure compared to physiological fluids (Bone & Denton, 1971, Hegele-Hartung et al., 1989), and an improper balance between the fixative and the cytoplasm can cause ultrastructural alterations where the intercellular space will be exaggerated due to hyper-osmolar fixatives or swelling of cells and organelles by hypo-osmolar fixatives (Bozzola & Russell, 1999). It may be possible that *Drosophila* neurons are more sensitive to osmotic pressures compared to other cellular targets, which could create ‘empty’ patches from cell lysis. It is also possible that these empty patches arise from subcellular regions that were washed out of the cell. Aldehyde-based fixatives are more selective for certain chemical groups over others, such as the amino acid lysine over carbohydrates and nucleic acids (Eltoum et al., 2001, McDonald & Auer, 2006). Unreacted macromolecules were most likely removed from the cells during the extensive washing, producing the empty patches that were sometimes seen. It is not well known if other artifacts, such as the disappearance/perturbation of microtubules, could arise due to the osmotic pressure imbalance between the fixative and the cytoplasm, or the chemical reactions of the fixatives themselves. Most past studies examining the effects of osmotic pressure and/or fixation on cellular ultrastructure have been done at a classical electron microscopy level (Bone & Denton, 1971, Bozzola & Russell, 1999, Hegele-Hartung et al., 1989). Such studies have many other steps post-fixation that could occlude observations, such as further fixation with osmium tetroxide, dehydration, resin-embedding, sectioning, and metal-staining. While chemical fixation has been incorporated into single-particle cryo-EM and cryo-ET, more studies will be needed to assess its impact on ultrastructural morphology.

All these efforts were instigated to address a potential bottleneck in the quality of samples for cryo-ET, an issue that has not been extensively talked about. Potential problems in cryo-EM involving the use of blotting for freezing samples has mostly been discussed and addressed at the single-particle level, with the development and implementation of specific instruments such as the Chameleon and the Vitrojet (Ravelli *et al*., 2020, Levitz *et al*., 2022). It’s possible that a similar innovation may need to be developed for the cellular level due to the issues highlighted here. Such issues, including blotting-induced varicosity formation that could eventually lead to cell death, is especially prominent in neurons, which possess mechanosensitive receptors and a large surface area by nature of its neurite network. To suppress artifactual varicosity formation in neurons while leaving physiologically relevant varicosities intact, new sample preparation techniques may need to be developed. There are several ways this could be implemented; one way is that the sample is frozen on an even faster timescale than plunge-freezing to minimize artifactual varicosity formation, which can happen on the order of seconds (Gu et al., 2017). High-pressure freezing could perhaps be one way around this, as freezing is typically achieved on the order of milliseconds (McDonald & Auer, 2006). Another method that could suppress varicosity artifacts is to co-culture the neurons with oligodendrocytes to confer protective myelination. This approach could be combined with micropatterning to create a model system for white matter, which are myelinated axons that are bundled closely together. Not only is this a more physiologically relevant system, but it is also more resistant to pressures by up to 39% compared to the unmyelinated grey matter (Budday *et al*., 2015). New tools and instruments, driven by the need to improve our models to be more physiologically relevant, will be necessary as we continue to investigate complex neurobiological questions that can be readily answered by the high-resolution capabilities offered by cryo-ET and its auxiliary techniques.

## MATERIAL AND METHODS

### Grid Preparation

Gold 200 mesh carbon or SiO_2_ R 2/1 grids or R 1.2/20 grids (Quantifoil Micro Tools GmbH) were coated with ∼8 nm of carbon using a Leica ACE600 (Leica Microsystems GmbH). The grids were glow-discharged either on a Harrick Plasma PDC32G115V Basic Plasma Cleaner (Harrick Plasma) or on a GloQube Plus (Quorum Technologies) to render them hydrophilic. Grids were then briefly cleaned with 70% ethanol (EtOH) followed by three washes in sterile, filtered water and one wash in Dulbecco’s phosphate buffered saline (DPBS, Lonza). For *Drosophila* neurons, grids were either placed in MatTek 35 mm/20 mm glass-bottom petri dishes (MatTek Life Sciences) and coated with sterile-filtered 0.5 mg/mL Concanavalin A, Alexa Fluor™ 350 (Invitrogen) in 1X phosphate-buffered saline (PBS, Corning Inc.) as the adherent extracellular matrix protein (ECM) or micropatterned (see following section). Grids were incubated overnight with the ECM in an incubator set at 25°C, 0% CO_2_ and then washed 3-5 times in H_2_O or 1X PBS before soaking in supplemented Schneider’s media (SSM), consisting of 80% Schneider’s media (Lonza) supplemented with 20% heat inactivated fetal bovine serum (FBS, ATCC) and 5 µg/mL insulin, 10 µg/mL tetracycline, 100 µg/mL streptomycin, and 100 µg/mL penicillin until the cell culturing step. For mouse neurons, grids were placed in MatTek 35 mm/20 mm glass-bottom petri dishes and coated with 0.1 mg/mL poly-D-Lysine (Gibco) and incubated overnight in an incubator set at 37°C, 5% CO_2_. Grids were then washed 3-5 times in H_2_O and then left in serum-free media (SFM), consisting of Neurobasal medium (Invitrogen) with B27 supplement (Invitrogen), 2 mM glutamine, 37.5 mM NaCl, and 0.3% glucose until the cell culturing step.

### Micropatterning

Micropatterning was performed as previously described (Sibert et al., 2021), with some modifications. Glow-discharged grids were placed onto pre-cleaned microscope glass slides, one grid per slide. Polydimethylsiloxane (PDMS) stencils (Alvéole Lab) were carefully placed over each grid with tweezers to secure them at the edges while minimizing contact with the carbon foil. The grids were then incubated in a humid chamber for at least 30 minutes in 10 µL of 0.05% poly-L-Lysine (Sigma-Aldrich), followed by 3-5 washes in filtered 0.1 M HEPES (pH 8.5). The grids were then incubated for at least 1 hour in 15 µL of 100 mg/mL polyethylene glycol-succinimidyl valerate (Laysan Bio, Inc.) mixed in 0.1 M HEPES as an antifouling layer. Grids were then washed five times with filtered H_2_O. The water was then aspirated from the grids until only a thin layer of liquid remained at the center of the grid. 4-benzoylbenzyl-trimethylammonium chloride gel (PLPP gel, Alvéole Lab) was then applied to the grid as a photocatalyst, either as a 1 µL volume alone or as a 5 µL solution of 0.8 µL gel mixed with 4.2 µL of 70% EtOH. The grids were then incubated for at least 20 minutes in the dark at room temperature until completely dry.

Grids were micropatterned using the PRIMO micropatterning system (Alvéole Lab) at a 30 mJ/mm^2^ dose and 100% laser power through the Leonardo software system. The grids were patterned using a curved or an X pattern, created through Adobe Illustrator and exported to the Leonardo software system as an uncompressed 8-bit .TIF file. 10 µL of sterile 1X PBS was immediately applied to the grids after the patterning finished for at least 10 minutes to dissolve any remaining PLPP gel. The stencil was then removed with a pair of tweezers, and the grids were sterilized with 15 µL of 70% EtOH for 2 minutes before washing the grids 5 times with sterile-filtered 1X PBS in 15 µL volumes. Grids were then placed in a 35 mm/20 mm dish filled with 1 mL of sterile filtered 0.5 mg/mL Concanavalin A, Alexa Fluor™ 350 in PBS as the adherent extracellular matrix protein to incubate overnight in an incubator at 25°C, 0% CO_2_. The micropatterned grids were then washed 3-5 times in H_2_O or PBS before being left in supplemented Schneider’s media (SSM) until the cell culturing step.

### Primary Cell Culture

*Drosophila* neurons were prepared as previously described with minor modifications (Lu et al., 2015, Egger et al., 2013, Sibert et al., 2021). Neurons were isolated from the following *Drosophila* genotypes: *elav-Gal4; UAS-mCD8::GFP* (Bloomington Drosophila Stock Center #5146), *elav-Gal4; UAS-Tau::GFP; UAS-DenMark* (*UAS-Tau::GFP* and *UAS-DenMark* correspond to Bloomington Stock Center #33052 and #33061, respectively). 30-40 3^rd^ instar larvae were isolated and screened for neuronal expression of mCD8::GFP on a Olympus SZX12 stereo microscope in fluorescence and brightfield mode. For the mCD8::GFP stock, larvae with fluorescently labeled neurons were easily identified by bright green fluorescence that illuminated the brain. To distinguish between dendrites and axons, we used a fly strain that expresses both the axon marker Tau::GFP and the dendrite marker DenMark(Nicolai *et al*., 2010) and has the Tubby(+) balancer, which causes larvae to have a short stout body and was used to eliminate flies that do not express these fluorescent markers. Positively identified larvae were then washed twice with 1X PBS, washed twice with 70% EtOH for sterilization, and washed twice with 1X sterile dissection saline consisting of 9.9 mM HEPES pH 7.5, 137 mM NaCl, 5.4 mM KCl, 0.17 mM NaH_2_PO_4_, 0.22 mM KH_2_PO_4_, 3.3 mM glucose, and 43.8 mM sucrose. Brains were individually isolated from larvae on the Olympus SZX12 stereo microscope in brightfield mode by first roughly tearing a larva at its midsection or head with a pair of sharp tweezers in a Sylgard® dissection dish filled with room-temperature 1X dissection saline. The isolated brains were stored in a tube of 1X dissection saline at room temperature until all brains were extracted. Brains were spun down for 1 minute at 300 x g and washed with 1X dissection saline twice. Supernatant was partially removed, and the brains were digested with 2.5 mg/mL Liberase™ (Thermolysin Medium) Research Grade (Roche) in 1X dissection saline and placed on a tube revolver rotator (ThermoFisher Scientific) for one hour. Brains were further digested by mechanical disruption while rotating by pipetting the solution 30 times every ten minutes with a P200 pipette to produce a cloudy suspension with body segments left over. The suspension was spun down for 5 minutes at 300 x g, and the digestion was stopped by adding SSM and pipetting 30 times to mix. The suspension was spun down again for 5 minutes at 300 x g, washed with SSM, and then filtered through a Falcon® 40 µm cell strainer (Corning Inc.) to remove the body segments from the cell suspension. The suspension was then spun down one final time and resuspended in 300-500 µL of SSM as the final cell suspension. The cell suspension was aliquoted in equal volumes onto plates with incubated grids for 30 minutes before flooding the plates with 2 mL of SSM. The neurons were monitored and grown for 3-4 days in an incubator set at 25°C, 0% CO_2_, where neurites were seen extending from the soma.

Cortical mouse neurons were cultured in accordance with NIH guidelines and approved by the University of Wisconsin Committee on Animal Care. Cortical (E15.5) neuron cultures were prepared from Swiss Webster mice as previously described (Viesselmann et al., 2011). Briefly, pregnant female mice were euthanized with CO_2_ and dissected to remove the uterus. Fetuses were decapitated into dissection medium (DM) consisting of Hank’s Balanced Salt Solution (Invitrogen) and HEPES. Brains were removed from the heads and dissected to remove the meninges and dissect the cortices in cold DM. Cortices were then digested with 2.5% trypsin and washed with plating medium (PM), consisting of Neurobasal medium with 5% FBS (Hyclone), B27 supplement, 2 mM glutamine, 37.5 mM NaCl and 0.3% glucose. Cortices were further digested by mechanical disruption by triturating with a P1000 pipette. Dissociated cortical neurons were then counted and plated onto plates with incubated grids at a density of 20,000 cells/cm^2^ for 1 hour in PM. After 1 hour, this medium was replaced with serum-free medium (SFM), which is PM without the FBS. The neurons were monitored and grown for up to 12 days in an incubator set at 37°C, 5% CO_2_, where neurites can be clearly seen emerging from the soma.

### Drug Treatment

Stock solutions of Paclitaxel (Cell Signaling Technologies) was created at 1 mM concentration by adding 1.15 mL of DMSO to powdered Paclitaxel, aliquoted, and stored at −80°C until further use. Subsequent dilutions of Paclitaxel were done in H_2_O to avoid possible cytotoxic effects from DMSO. Before adding to the culture, Paclitaxel was diluted to two times the target concentration in SSM, and then added to the culture by aspirating 1 mL of SSM from the culture and then adding 1 mL of the Paclitaxel + SSM solution to the final concentration of 5 or 20 nM. The neurons were cultured for at least 24 hours before freezing.

### Fluorescence Imaging

Before plunge-freezing, neurons were imaged on a Leica DMi8 widefield fluorescence (Leica Microsystems GmbH) microscope at room temperature. To assess culture quality, the entire grid was imaged at 40X magnification (0.6 NA air objective) in brightfield, GFP (emission, λ = 525 nm), and DAPI (emission, λ = 477 nm) channels using the Leica LAS X software in navigator mode and stitched together. To assess neuronal quality for the blotting tests, images were acquired at 100X magnification (1.4 NA oil objective) before and after blotting at the center of the grid for easier identification and orientation.

### Fixation

Half of the culture media was aspirated from the cell culture. Neurons were then fixed for 10-20 minutes in 4% paraformaldehyde (Methanol-free, EM grade, Electron Microscopy Sciences) in Krebs buffer and sucrose (PKS) (Dent & Meiri, 1992) containing 145 mM NaCl, 5 mM KCl, 1.2 mM CaCl, 1.3 mM MgCl_2_, 1.2 mM NaH_2_PO_4_, 10 mM glucose, 20 mM HEPES, and 0.4 mM sucrose, 4% PKS with 0.25% glutaraldehyde (EM grade, Ted Paella), 4% paraformaldehyde in cytoskeleton buffer containing 80 mM PIPES pH 6.9, 2 mM MgCl_2_, and 0.5 mM EGTA (Cytoskeleton Inc.), 2.5% glutaraldehyde in 0.1 M sodium cacodylate buffer pH 7.4 (Electron Microscopy Sciences), or 2.5% glutaraldehyde in 0.1 M Sodium Cacodylate buffer pH 7.4 with 2% paraformaldehyde in 1 mL volumes. Neurons were then carefully washed with 2 mL of sterile 1X PBS or 0.1 M sodium cacodylate buffer 10 times at 5-minute intervals before freezing. Neurons that were not fixed were still washed with sterile 1X PBS at the same volume and interval before freezing.

### Osmolality measurements

All osmolality measurements were done carried out with a Type 6/6M Micro-osmometer in accordance to manufacturer protocol using freezing point depression at 100 µL volumes per measurement (Löser Messtechnik). An initial calibration was carried out with solutions of dI H_2_O (0 mOsm) and two solutions of NaCl (300 mOsm and 900 mOsm respectively). Osmolality measurements were then carried out in triplicates with 26 internal standards of KCl solutions with a molality range from 0.311 to 0.8618, giving a range of osmolalities from 58 to 1491 (in miliosmoles). These measured osmolalities were then adjusted with a Platford correction factor (PCF) using osmolalities of KCl measured at 0°C (Platford, 1973). The PCF was plotted against the KCl osmolalities to create a calibration curve fitted with a 4^th^ degree polynomial. Raw osmolality values were measured with the sample solutions (Table 4) in triplicates, averaged, and then corrected by the calibration curve to give the final osmolalites. Out of the 11 samples tested, two solutions (4% PKS and 4% PKS with 0.25% GA) gave raw osmolality values that were too high for the calibration curve to accurately correct and are hence denoted as rough estimates.

### Blotting and Freezing

Grids were blotted and plunge-frozen using the Leica EM GP (Leica Microsystems GmbH) or the Gatan CP3 (Gatan, Inc.). Grids were picked up from the glass bottom MatTek dish using 5/15 Style Dumont Tweezers (Electron Microscopy Sciences), transferred to their respective tweezers and placed into the EM GP chamber set at 25°C and 99% humidity or the CP3 chamber. For freezing, 4 µL of pre-warmed 10 nm gold fiducials (Aurion Gold Nanoparticles, Electron Microscopy Sciences) was added to the grids, blotted for 6-12 seconds from the backside of the grid, and then plunge-frozen into liquid ethane cooled by liquid nitrogen. For the blotting test, the grid was blotted at the same settings as the freezing parameters, but then placed back into the cell culture dish without plunging into liquid ethane by using the ‘Test blot’ feature on the EM GP interface and then immediately imaged on the DMi8 widefield fluorescent microscope afterwards. Frozen grids were stored in liquid nitrogen until data acquisition.

### Cryo-ET

Plunge-frozen grids were clipped into autogrids and loaded into a Talos Arctica 200 kV transmission electron microscope for imaging or Titan Krios 300 kV transmission electron microscope for imaging and tilt series data collection (ThermoFisher Scientific). Images were acquired with a Falcon3 direct-electron detector in linear mode (ThermoFisher Scientific) or a K3 direct-electron detector in EFTEM mode with a BioQuantum energy filter set at 20 eV slit width (Gatan Inc.). Atlases for each grid were collected at a low magnification of 82x (1,101 Å/pixel or 1,097 Å/pixel after microscope upgrade and recalibration for fringe-free imaging) or 110x (773 Å/pixel). Overview cryo-EM maps were taken at a magnification range of 2,850x – 6,500x (28.5 – 13.7 Å/pixel). From the overview cryo-EM maps, cellular regions worthy of tilt series data collection were identified by the presence of intact neurites.

Single tilt series on mouse cortical neurons and *Drosophila* neurons fixed in PKS + GA were collected from −60° to 60° in 2-3° increments in a bidirectional or a dose-symmetric collection scheme (Hagen *et al*., 2017) in CDS counting mode on the K3 with SerialEM (Mastronarde, 2005) at 15,000x (pixel size 6.15 Å/pixel) or 19,500x magnification (pixel size 4.603 Å/pixel) at a nominal defocus of 8 µm at a total dose of 85-100 e^-^/Å^2^. Frames were motion-corrected by MotionCor2 (Zheng *et al*., 2017). Tilt series were aligned by tracking 10 nm gold fiducial particles, binned by 2 to a pixel size of 9.206 Å/pixel, CTF-corrected with ctfplotter and ctfphaseflip (Xiong *et al*., 2009), and reconstructed into tomograms by weighted back projection in IMOD (Kremer *et al*., 1996). Tomograms were low-pass filtered to 80 Å using EMAN2 (Tang *et al*., 2007) and examined in IMOD. Segmentation was done using the convoluted neural network in EMAN2 (Chen *et al*., 2017) and viewed on Chimera or ChimeraX (Pettersen *et al*., 2004, Pettersen *et al*., 2021) Montage tilt series in a 3×3 or a 3×4 tilt pattern, with an overlap of 10-20% in X and 10% in Y were collected from −60° to 60° in 3° increments at 19,500x magnification (pixel size 4.603 Å/pixel or 4.517 Å/pixel after microscope upgrade and recalibration for fringe-free imaging) in a dose-symmetric collection scheme in CDS counting mode with SerialEM through a custom script at a total dose of 50-75 e^-^/Å^2^ (https://github.com/wright-cemrc-projects/cryoet-montage/tree/main/SerialEM) with default spiral translation parameters and a nominal defocus of 3-8 µm. Frames were motion-corrected by MotionCor2.

Montage tilt series were stitched together by a custom python script (https://github.com/wright-cemrc-projects/cryoet-montage/tree/main/Python), with occasional correction using the manual image alignment package in IMOD. Fully stitched montage tilt series were then binned by 2 to a pixel size of 9.206 or 9.034 Å/pixel, aligned with gold fiducials or patch tracking (if gold fiducials were not present), CTF corrected with ctfplotter and ctfphaseflip, and reconstructed by weighted back-projection in IMOD. Tomograms were further binned by 2 to a final pixel size of 18.4 or 18.1 Å/pixel, and then low-pass filtered to 80 Å using EMAN2 and examined on IMOD. Segmentation was done using the convoluted neural network in EMAN2 and viewed on Chimera or ChimeraX.

### Varicosity Measurements

Measurements of varicosity diameter and its adjacent shafts were done on IMOD. Overview cryo-EM maps for four sample conditions A-D were collected and stitched together by blendmont in IMOD (Table 1). In accordance with a previous study, a varicosity was defined when an oblong-shaped swelling along the neurite was ≥ 300% than the adjacent neurite shaft (Ma et al., 2022). For each varicosity, the diameter was measured (D_Varicosity_) at the midpoint between the two ends, defined by when the swelling tapers off to a neurite shaft with the membranes parallel to each other. For each varicosity, four additional diameters were measured at the adjacent neurite shaft; two diameters at the immediate ends of the swelling where the membrane becomes parallel with each other (D_Shaft1A_ and D_Shaft1B_), and two diameters 250 nm away from D_Shaft1A_ and D_Shaft1B_ at the opposite direction from the varicosity center (D_Shaft2A_ and D_Shaft2B_). D_Shaft1A_ and D_Shaft1B_ were averaged to a final ‘Adjacent’ D_Shaft1_, and D_Shaft2A_ and D_Shaft2B_ were averaged to a final ‘Away’ D_Shaft2_. In total, measurements were collected from 96 varicosities in 41 maps from sample condition A, 95 varicosities in 29 maps from sample condition B, 93 varicosities in 19 overview maps from sample condition C, and 95 varicosities in 5 maps from sample condition D. The measurements were analyzed with Prism 9 (GraphPad).

---

**Supplemental Figure 1.**
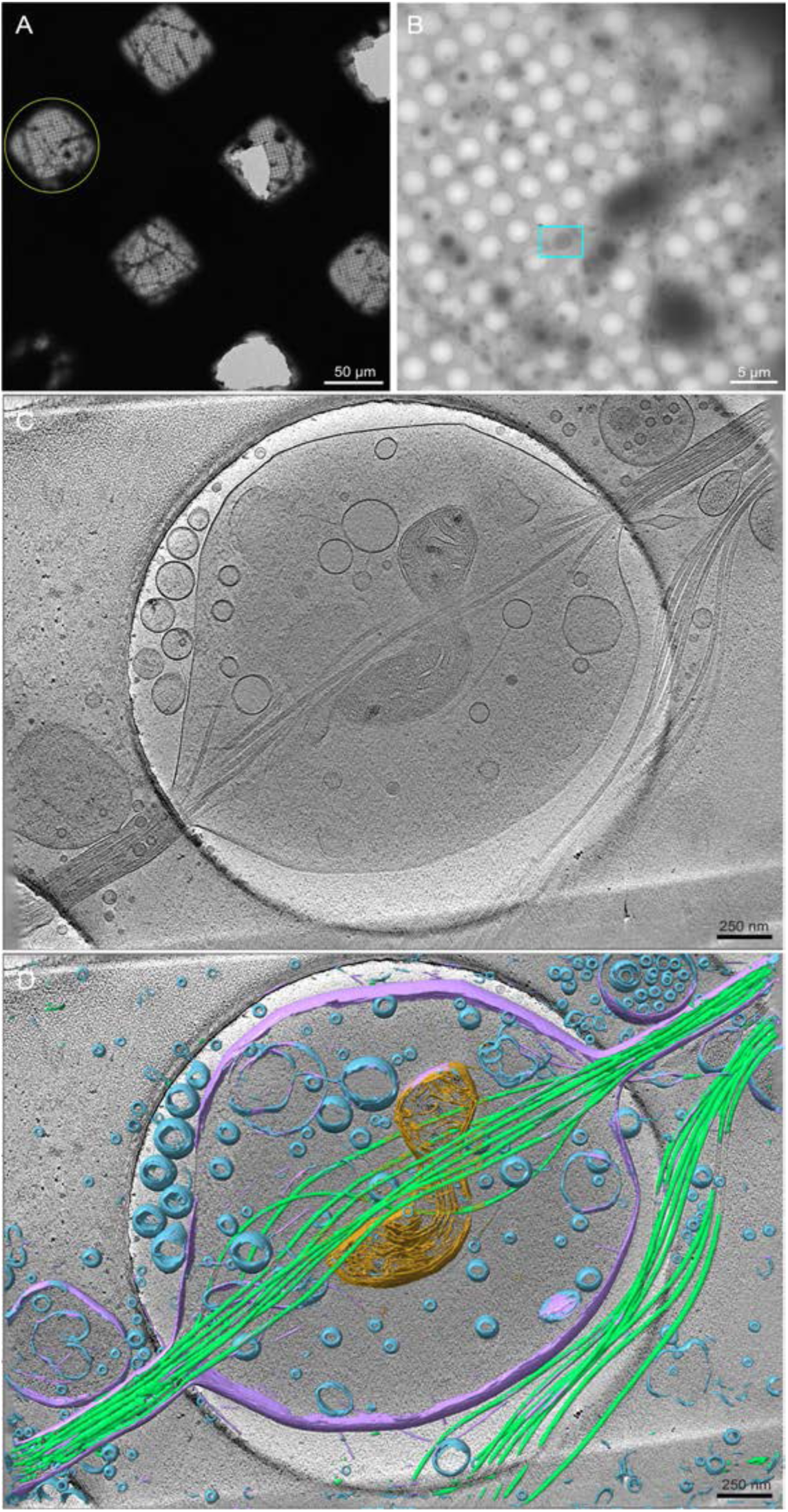
Cryo-EM and cryo-ET of primary cortical neurons derived from *Mus musculus*, grown for 12 days *in vitro* (DIV). **(A)** Low magnification cryo-EM image montage of mouse neurons, frozen with a Gatan CP3. **(B)** Magnified cryo-EM grid square montage of the yellow circle in (A). **(C)** A 50 nm thick tomogram slice from a tilt series that was taken at the cyan-marked box in (B). **(D)** Segmentation of the tomogram by a convolutional neural network in EMAN2, overlaid on top of the 50 nm slice in (C). Peripheral membranes and membrane blebs are colored in purple, microtubules are colored in light green, vesicles are colored in blue, and a mitochondrion is colored in orange. The scale bars in (A)-(D) are embedded in the image. Pixel sizes for (A) is 1,101 Å/pixel (82x magnification), 17.1 Å/pixel (4,800x magnification) for (C), and 6.15 Å/pixel (15,000x magnification) for (D) and (E).

**Supplemental Figure 2.**
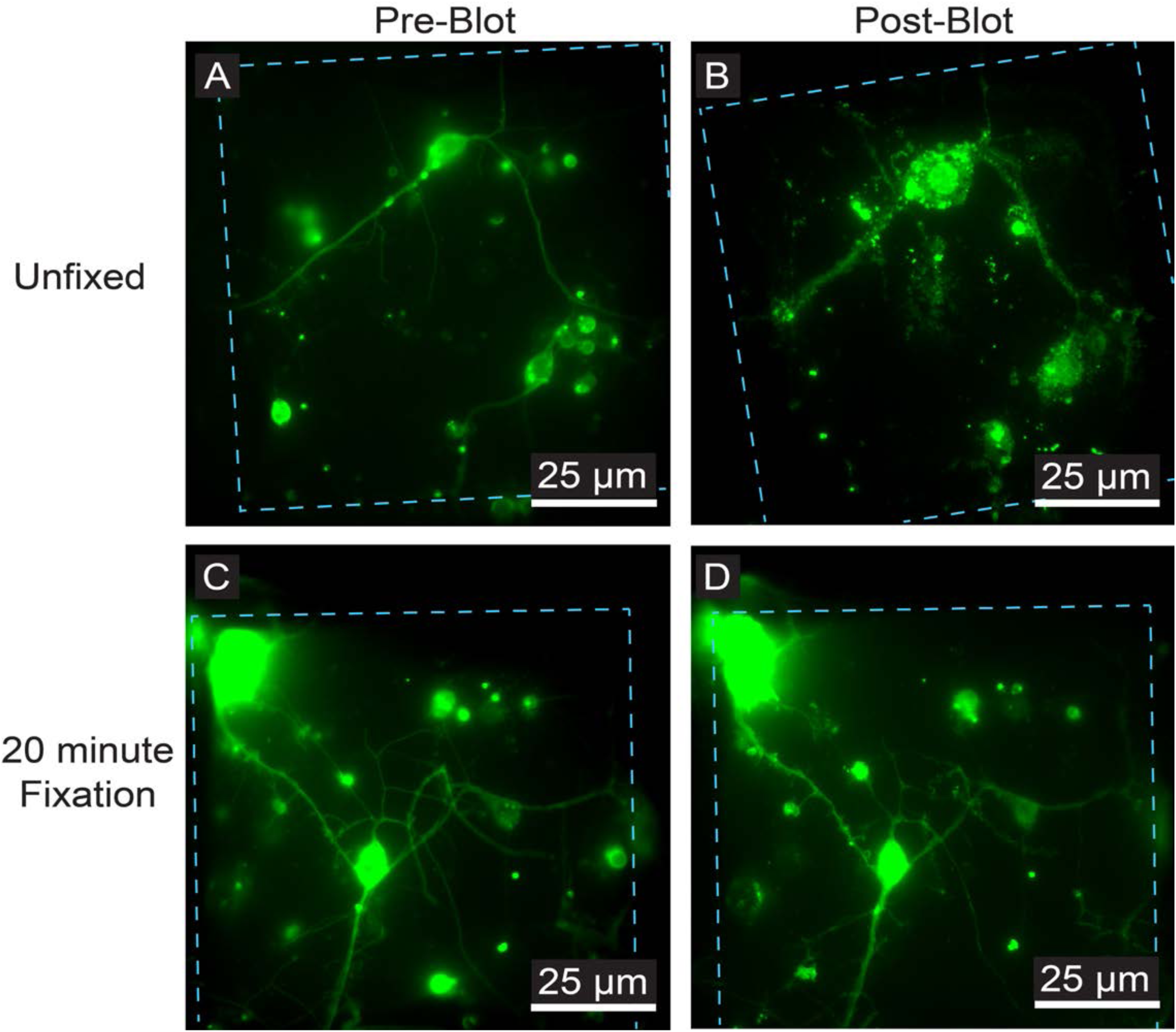
Effects of blotting on primary *Drosophila* neurons cultured on unpatterned EM grids under regular or chemical fixation conditions. **(A)** Live-cell fluorescence microscopy (FLM) of a *Drosophila* neuron expressing membrane-targeted GFP cultured on unpatterned R 1.2/20 grid squares coated with concanavalin A. **(B)** The same square in (A), re-imaged after filter paper blotting the grid using a Leica EM GP. Note the loss of morphological integrity at both the cell body and the neurites, seen by diffuse green fluorescence along the neurites and cell body accompanied by swelling. Finer secondary neurite features are also lost. **(C)** FLM image of a *Drosophila* neuron expressing membrane-targeted GFP cultured on unpatterned R 1.2/20 grid squares coated with concanavalin A. **(D)** The same square in (C), re-imaged after chemical fixation in 4% PKS for 20 minutes and then filter paper blotting the grid using a Leica EM GP. Unlike (B), a vast majority of the cell remains intact after blotting, including fine secondary neurites branching off the primary neurites. Grid bars are represented by the cyan dashed line. The scale bars in (A-D) are embedded in the image. Green: *Drosophila* neurons.

**Supplemental Figure 3.**
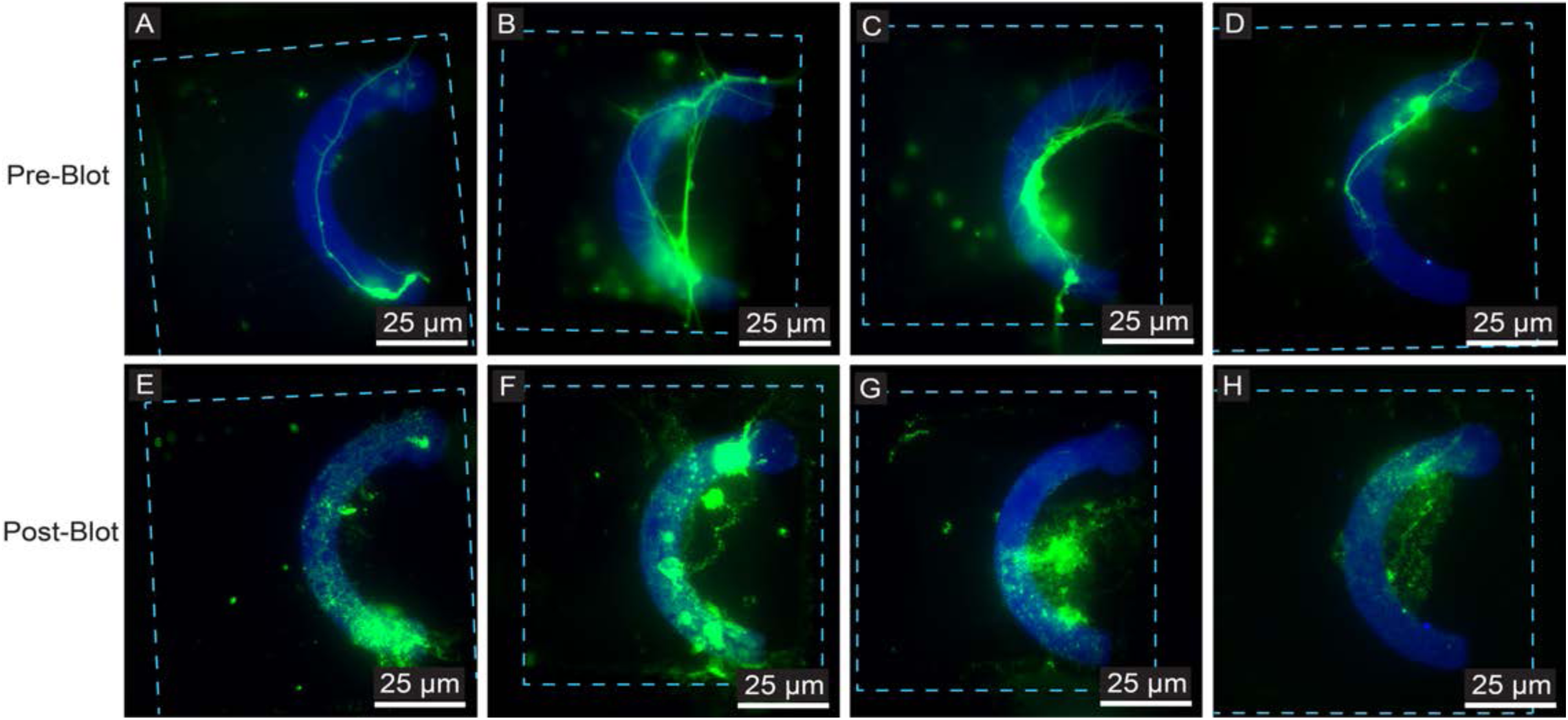
Examples of blotting effects on primary *Drosophila* neurons that were cultured on micropatterned EM grids under regular conditions. **(A-D)** Four representative images of live-cell fluorescence microscopy (FLM) of *Drosophila* neurons expressing membrane-targeted GFP cultured on micropatterned R 1.2/20 EM grid squares with fluorescent concanavalin A. **(E-H)** The same squares in (A-D) respectively, re-imaged after filter paper blotting the grid on a Leica EM GP. Note the loss of morphological integrity at both the cell body and the neurites, seen by diffuse green fluorescence along the neurites and cell body, loss of finer secondary neurites, and swelling along the cell body and the neurites. Grid bars are represented by the cyan dashed line. The scale bars in (A-H) are embedded in the image. Green: *Drosophila* neurons. Blue: Micropattern.

**Supplemental Figure 4.**
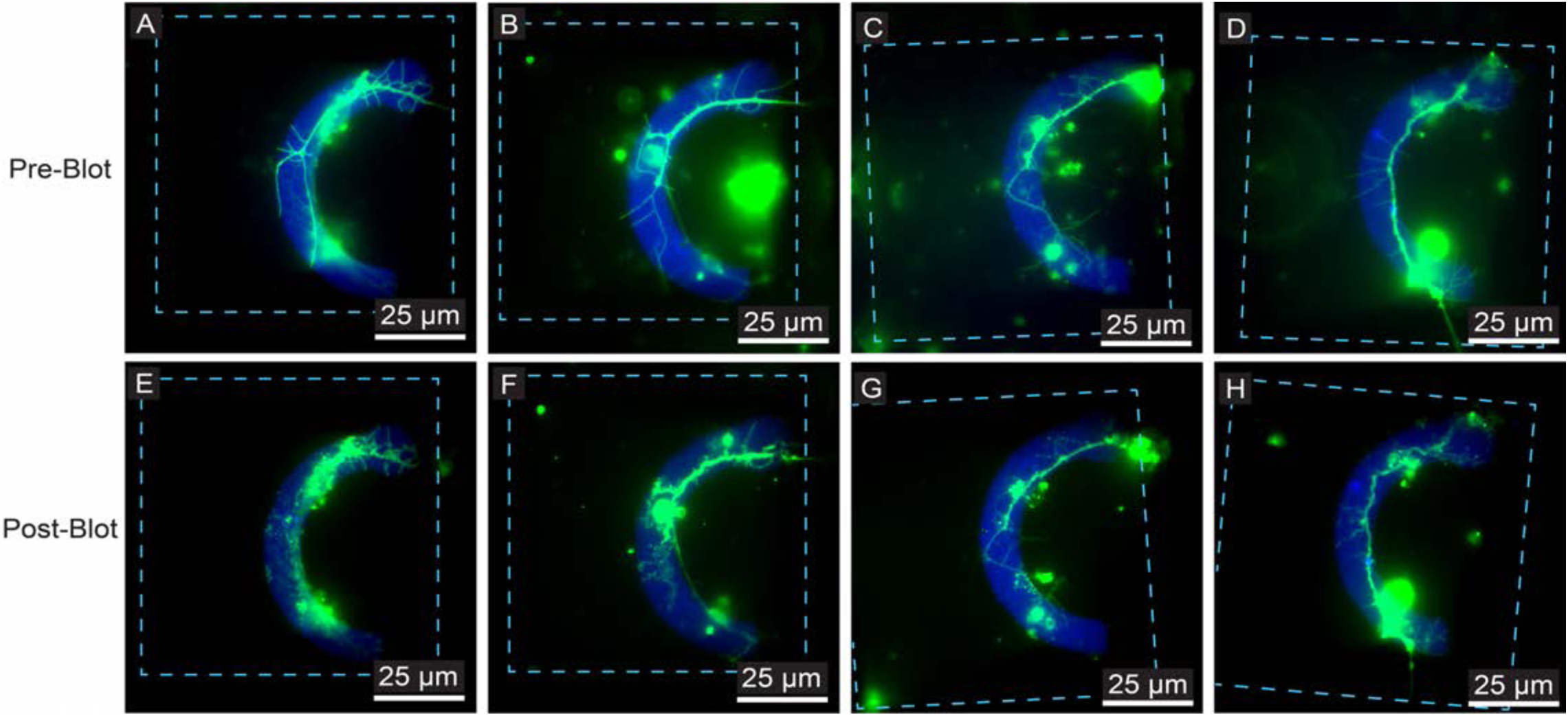
Examples of blotting effects on primary *Drosophila* neurons cultured on micropatterned EM grids with chemical fixation. **(A-D)** Four representative images of live-cell fluorescence microscopy (FLM) of a *Drosophila* neuron expressing membrane-targeted GFP cultured on micropatterned R 1.2/20 EM grid squares with fluorescent concanavalin A. **(E-H)** The same squares in (A-D) respectively, re-imaged after chemically fixing the grid in 4% PKS for 20 minutes and then filter paper blotting the grid on a Leica EM GP. The vast majority of the fixed cells in each square remain intact after blotting, including fine secondary neurites branching off the primary neurites. Grid bars are represented by the cyan dashed line. The scale bars in (A-H) are embedded in the image. Green: *Drosophila* neurons. Blue: Micropattern.

**Supplemental Figure 5.**
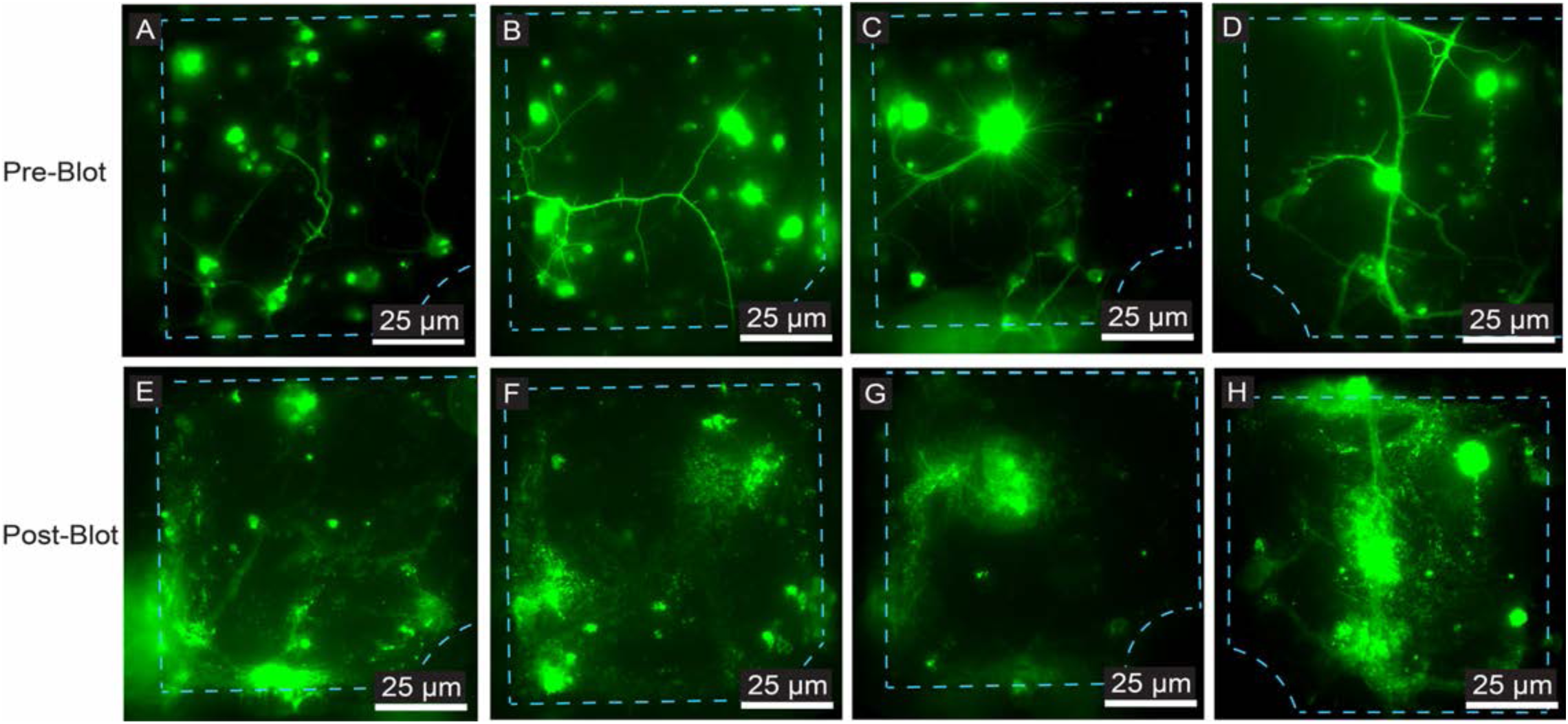
Examples of blotting effects on primary *Drosophila* neurons cultured on unpatterned EM grids under regular conditions. **(A-D)** Four representative live-cell FLM images of *Drosophila* neurons expressing membrane-targeted GFP cultured on unpatterned R 1.2/20 EM grid squares coated with concanavalin A. **(E-H)** The same squares in (A-D) respectively, re-imaged after blotting the grid on the Leica EM GP. Note the loss of morphological integrity at both the cell body and the neurites, seen by diffuse green fluorescence along the neurites and cell body, loss of finer secondary neurites, and swelling along the cell body and the neurites. Grid bars are represented by the cyan dashed line. The scale bars in (A-H) are embedded in the image. Green: *Drosophila* neurons.

**Supplemental Figure 6.**
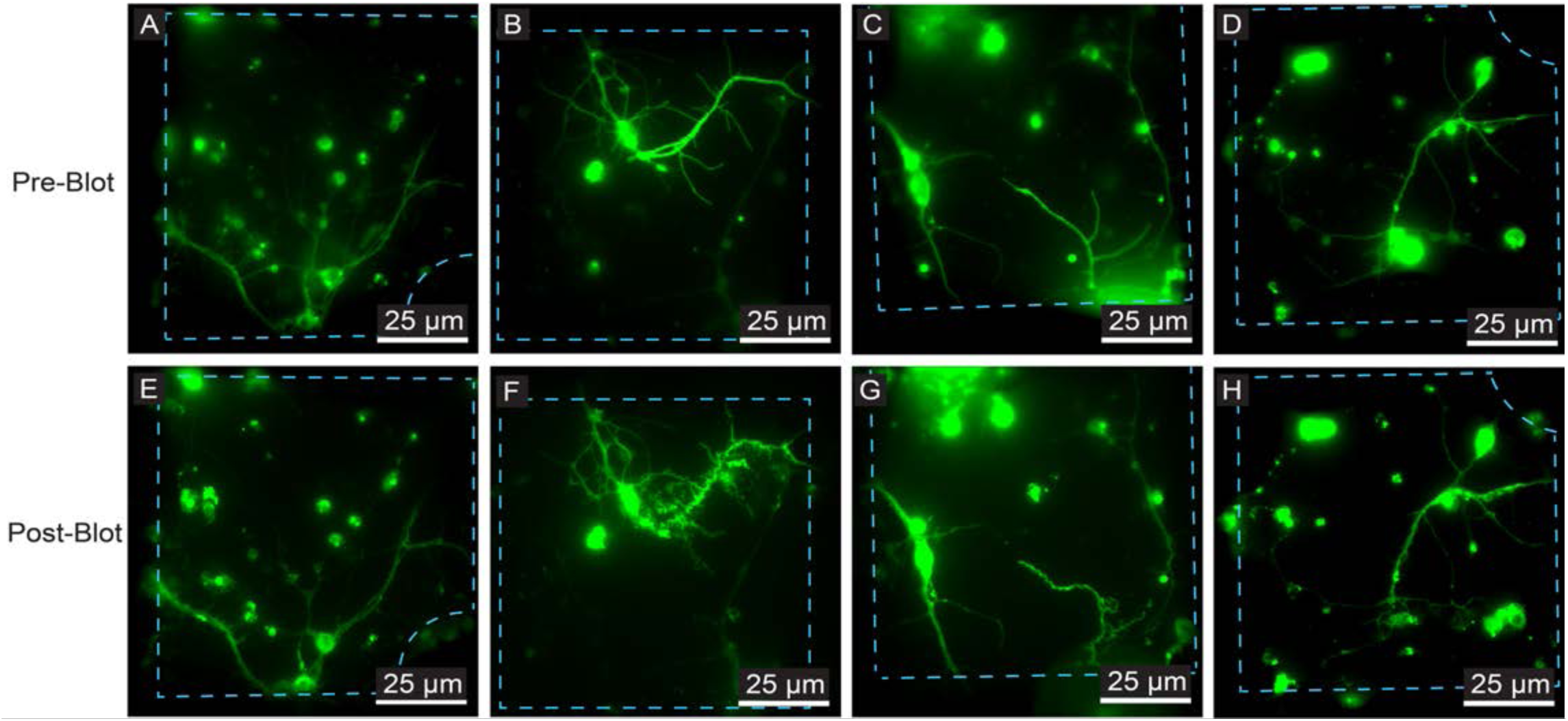
Examples of blotting effects on primary *Drosophila* neurons cultured on unpatterned EM grids with chemical fixation. **(A-D)** Four representative live-cell FLM images of *Drosophila* neurons expressing membrane-targeted GFP cultured on unpatterned R 1.2/20 EM grid squares coated with concanavalin A. **(E-H)** The same squares in (A-D) respectively, re-imaged after chemically fixing the grid in 4% PKS for 20 minutes and then blotting the grid on the EM GP. The vast majority of the fixed cells in each square remains intact after blotting, including fine secondary neurites branching off the primary neurites. Grid bars are represented by the cyan dashed line. The scale bars in (A-H) are embedded in the image. Green: *Drosophila* neurons.

**Supplemental Figure 7.**
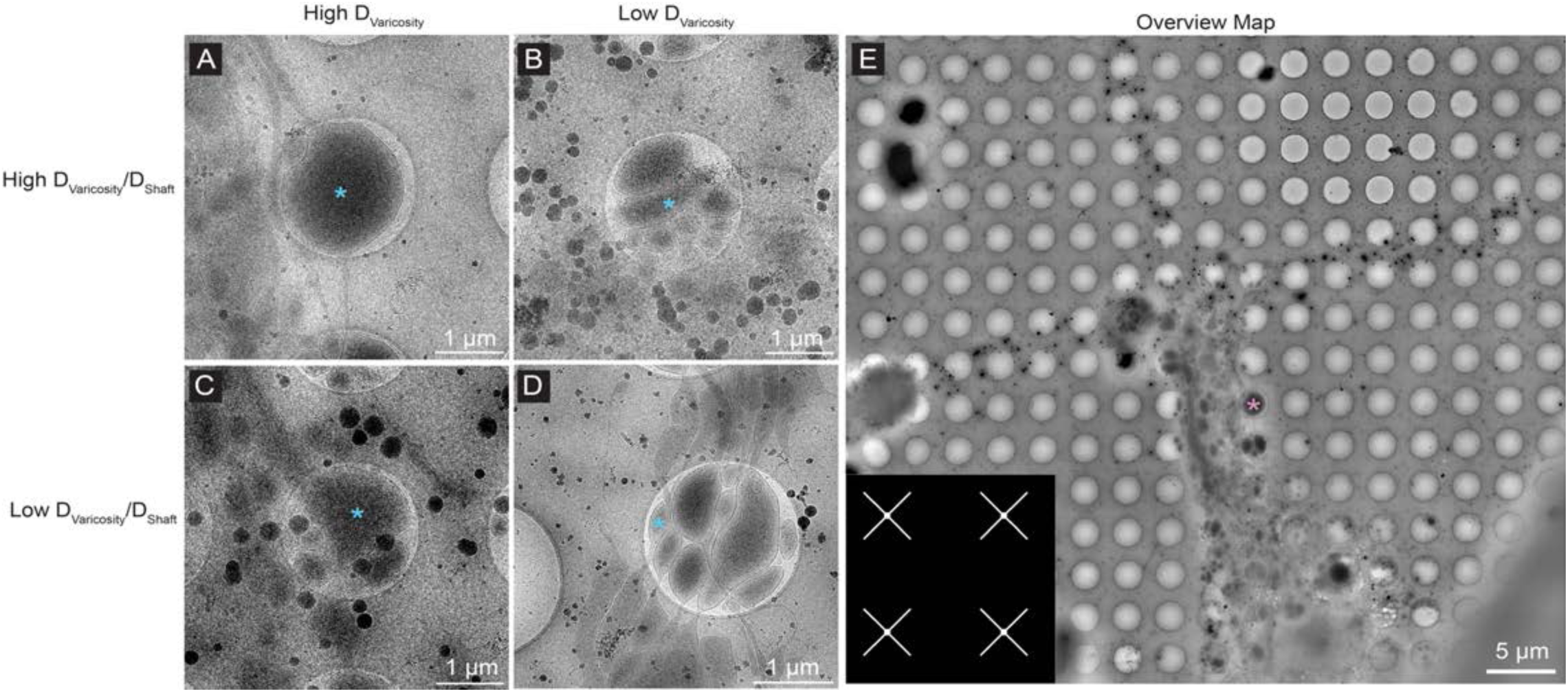
Categorization of varicosities based on the diameters of the varicosities (D_Varicosity_) and the ratio between D_Varicosity_ and D_Shaft_ from unfixed *Drosophila* neurons on X-patterned R 2/1 grids. **(A)** An unfixed varicosity class defined by having a high D_Varicosity_ and a high D_Varicosity_:D_Shaft_ ratio. **(B)** A class defined by having a low D_Varicosity_ and a high D_Varicosity_:D_Shaft_ ratio. **(C)** A class defined by having a high D_Varicosity_ and a low D_Varicosity_:D_Shaft_ ratio. **(D)** A class defined by having a low D_Varicosity_ and a low D_Varicosity_:D_Shaft_ ratio. **(E)** An overview cryo-EM map of a *Drosophila* neuron cultured on X-patterned R 2/1 grids, plunge-frozen on a Gatan CP3 without chemical fixation. The pattern used is shown in the inset, and the pink asterisk is the varicosity class in (A). The scale bars in (A-E) are embedded in the image. Pixel sizes are 28.5 Å/pixel (2,850x magnification) for (A-C) and (E), and 16.14 Å/pixel (5,000x magnification) for (D).

**Supplemental Figure 8.**
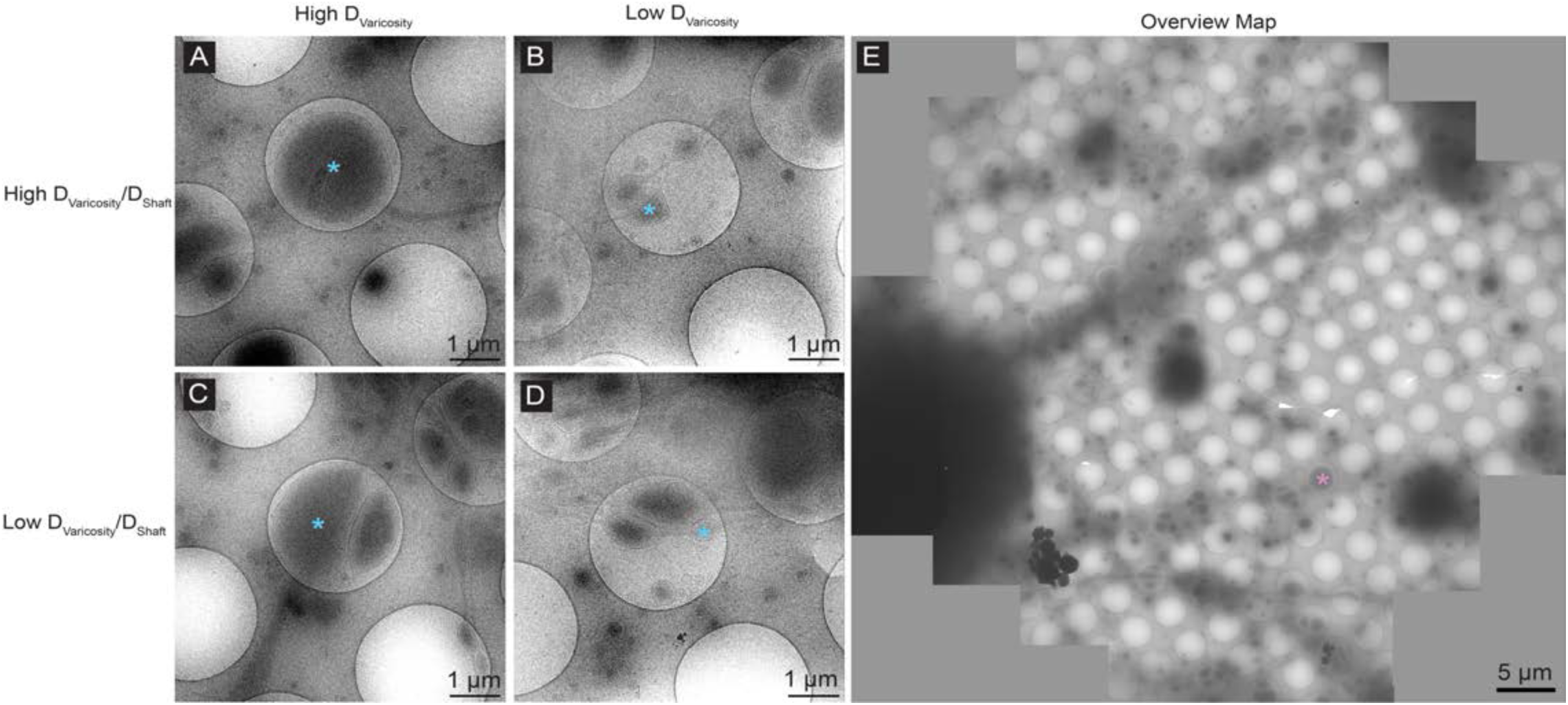
Categorization of varicosities based on the diameters of the varicosities (D_Varicosity_) and the ratio between D_Varicosity_ and D_Shaft_ from unfixed *Mus musculus* neurons on unpatterned R 2/1 grids. **(A)** An unfixed varicosity class defined by having a high D_Varicosity_ and a high D_Varicosity_:D_Shaft_ ratio. **(B)** A class defined by having a low D_Varicosity_ and a high D_Varicosity_:D_Shaft_ ratio. **(C)** A class defined by having a high D_Varicosity_ and a low D_Varicosity_:D_Shaft_ ratio. **(D)** A class defined by having a low D_Varicosity_ and a low D_Varicosity_:D_Shaft_ ratio. **(E)** An overview cryo-EM map of a *Mus musculus* neuron cultured on unpatterned R 2/1 grids, plunge-frozen on a Gatan CP3 without chemical fixation. The pink asterisk is the varicosity class in (A). The scale bars in (A-E) are embedded in the image. Pixel size is 17.1 Å/pixel (4,800x magnification) for (A-E).

**Supplemental Figure 9.**
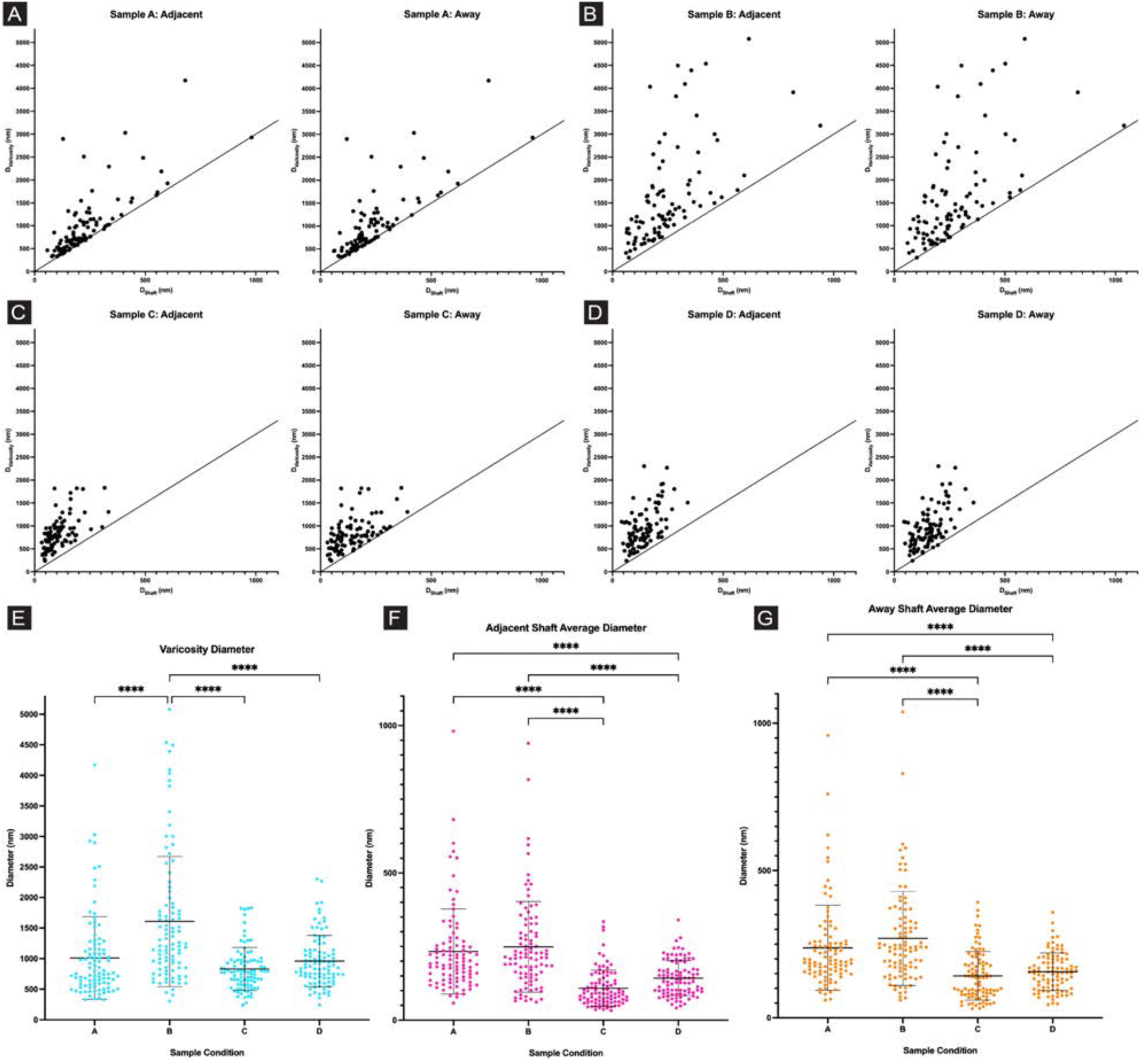
Measurements of D_Varicosity_, D_Shaft1_ (Adjacent), and D_Shaft2_ (Away). **(A)** Scatter plot of measurements from sample condition A (*Drosophila* neurons, curve-patterned R 1.2/20 grids, frozen with chemical fixation) with D_Varicosity_ plotted against D_Shaft1_ (left) and D_Shaft2_ (right). **(B)** Scatter plot of measurements from sample condition B (*Drosophila* neurons, curve-patterned R 1.2/20 grids, frozen without chemical fixation) with D_Varicosity_ plotted against D_Shaft1_ (left) and D_Shaft2_ (right). **(C)** Scatter plot of measurements from sample condition C (*Drosophila* neurons, X-patterned R 2/1 grids, frozen without chemical fixation) with D_Varicosity_ plotted against D_Shaft1_ (left) and D_Shaft2_ (right). (**D)** Scatter plot of measurements from sample condition D (*Mus musculus* neurons, unpatterned R 2/1 grids, frozen without chemical fixation) with D_Varicosity_ plotted against D_Shaft1_ (left) and D_Shaft2_ (right). **(E)** Column scatter plot of D_Varicosity_, colored cyan to match Fig 7A-D. **(F)** Column scatter plot of averaged D_Shaft1_, colored pink to match Fig 7A-D. **(G)** Column scatter plot of averaged D_Shaft2_, colored orange to match Fig 7A-D. N = 96 varicosities for sample condition A, N = 95 for B, N = 93 for C, N = 95 for D. All data are mean ± s.d. * is P < 0.05, ** is P < 0.01, *** is P < 0.001, **** is P < 0.0001 calculated using a one-way ANOVA with post-hoc Tukey’s multiple comparison test.

**Supplemental Figure 10.**
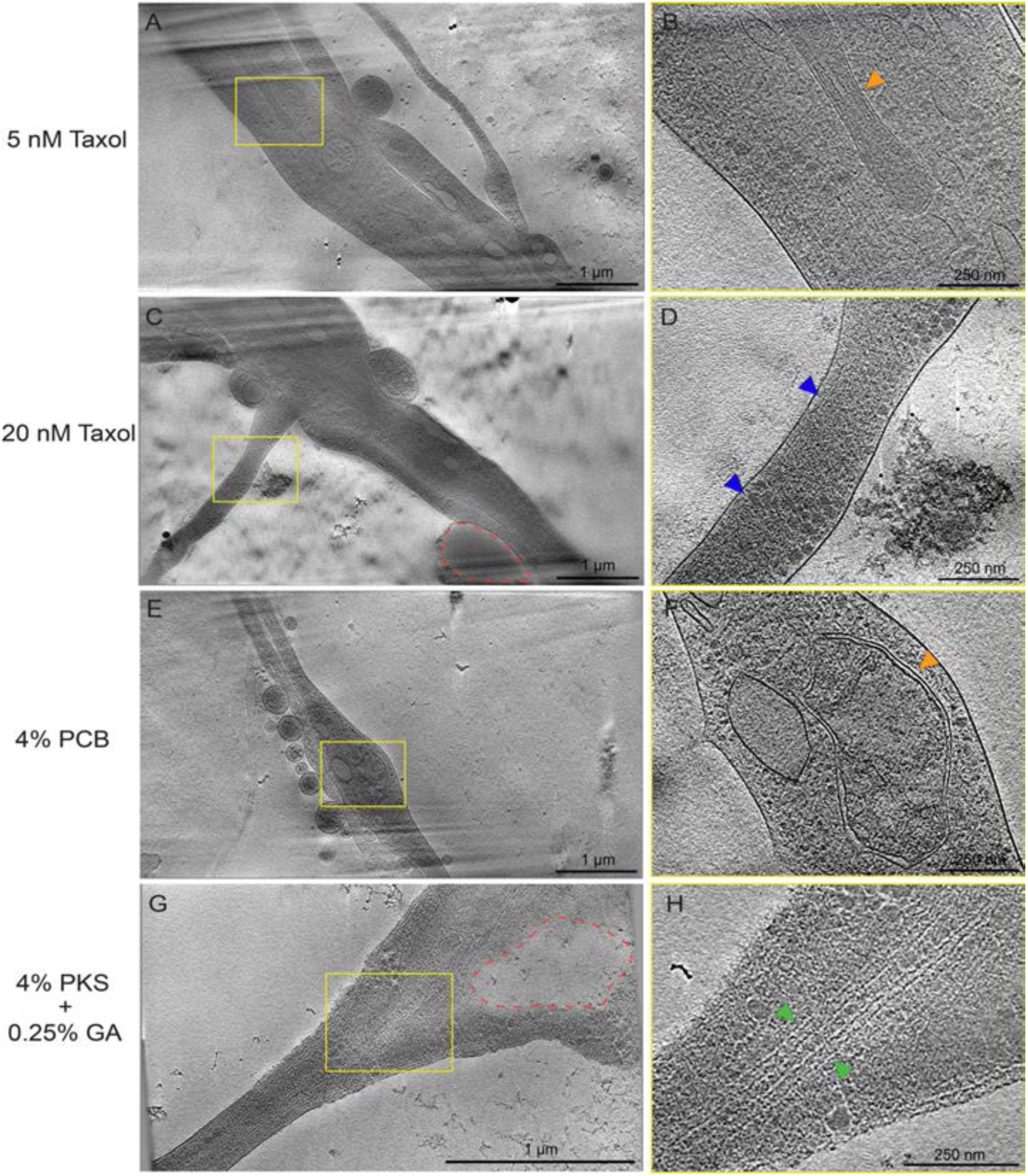
Attempts to mitigate disruption of microtubules by chemical fixation. (A) A 50 nm thick 3×3 montage tomogram slice on a Drosophila neuron cultured on curve-patterned R 1.2/20 grids, incubated with 5 nM of Taxol for 24 hours before chemical fixation and freezing. (B) A magnified view of a 50 nm thick tomogram slice at the yellow box highlighted in (A). Despite being able to see the mitochondria (orange arrowhead), microtubules were not found throughout the tomogram. (C) A 50 nm thick 3×3 montage tomogram slice on a Drosophila neuron cultured on curve-patterned R 1.2/20 grids, incubated with 20 nM of Taxol for 24 hours before chemical fixation and freezing. The red dotted line depicts a possible artifact where there is a sharp boundary between regions containing subcellular density and empty, or ‘cleared’ areas (D) A magnified view of a 50 nm thick tomogram slice at the yellow box highlighted in (C). Despite being able to see actin (blue arrowhead), microtubules were not found throughout the tomogram. (E) A 50 nm thick 3×3 montage tomogram slice on a Drosophila neuron cultured on curve-patterned R 1.2/20 grid, frozen after being chemically fixed in 4% paraformaldehyde dissolved in cytoskeleton buffer (PCB). (F) A magnified view of a 50 nm thick tomogram slice at the yellow box highlighted in (E). Despite being able to see mitochondria (orange arrowhead), microtubules were not found throughout the tomogram. (G) A 50 nm thick single tomogram slice on a Drosophila neuron cultured on an unpatterned R 1.2/20 grid, frozen after being chemically fixed in 4% PKS that also had 0.25% glutaraldehyde (GA) added. The red dotted line depicts a possible artifact where there is a sharp boundary between regions containing subcellular density and empty, or ‘cleared’ areas (H) A zoomed up view of a 25 nm thick tomogram slice at the yellow box highlighted in (G). Microtubules (green arrowhead) can be seen, however they are not at the quality of unfixed or GA-only fixed neurons that is seen in Fig 8. (I) A 50 nm thick 3×3 montage tomogram slice on a Drosophila neuron cultured on an curve-patterned R 1.2/20 grid, frozen after being chemically fixed in 2.5% GA and 2% PFA in cacodylate buffer. (J) A magnified view of a 40 nm thick tomogram slice at the yellow box highlighted in (I). Microtubules (green arrowhead) can be seen, however they are not at the quality of unfixed or GA-only fixed neurons that is seen in Fig 8. Scale bars are embedded in the image. (A-F) and (I-J) are at a pixel size of 4.518 Å/pixel (19,500x magnification), while (G-H) are at a pixel size of 4.603 Å/pixel (19,500x magnification).

## ACKNOWLEDGEMENTS

We would like to thank Dr. Rebeccah J. Katzenberger and Dr. David A. Wassarman in the Department of Genetics at the University of Wisconsin-Madison for additional help in raising and maintaining fly strains and crosses. We would also like to thank Emily Zytkiewicz and Dr. Thomas Record in the Department of Biochemistry at the University of Wisconsin-Madison for the use of their osmometer and the helpful discussions in osmolality. This work was supported in part by the University of Wisconsin-Madison, the Department of Biochemistry at the University of Wisconsin-Madison, and public health service grants R01 GM114561 and U24 GM139168 to E.R.W., R01NS102385 to J.W, and R01 NS115400 to E.W.D. from the NIH. L.A.E. was supported through the Science and Medicine Graduate Research Scholars Program at the University of Wisconsin-Madison. We are grateful for the use of facilities and instrumentation at the Cryo-EM Research Center in the Department of Biochemistry at the University of Wisconsin-Madison.

